## Supplementary Figures for "Identification and Characterization of Archaeal Pseudomurein Biosynthesis Genes through Pangenomics"

**(a)**

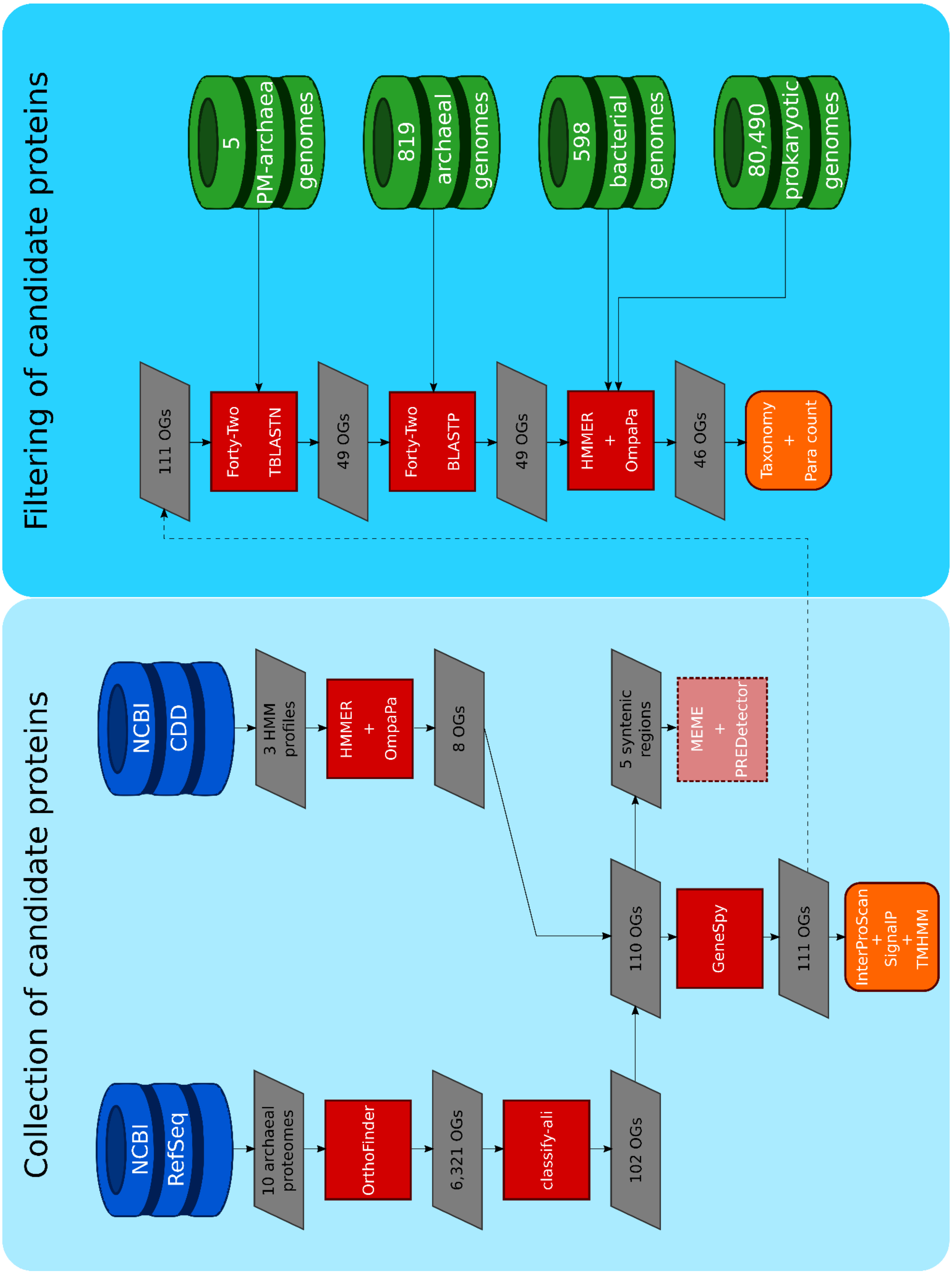


**Figure S1. Overview of the methods used for the bioinformatic analyses carried out during this study (see Material and Methods of the main text for**

**details). (a, previous page)** Main steps of the pipeline for the identification and filtering of the OGs potentially involved in PM biosynthesis. **(b, this page)** Details of the datasets and AA substitution models used for phylogenetic analyses.

**(b)**

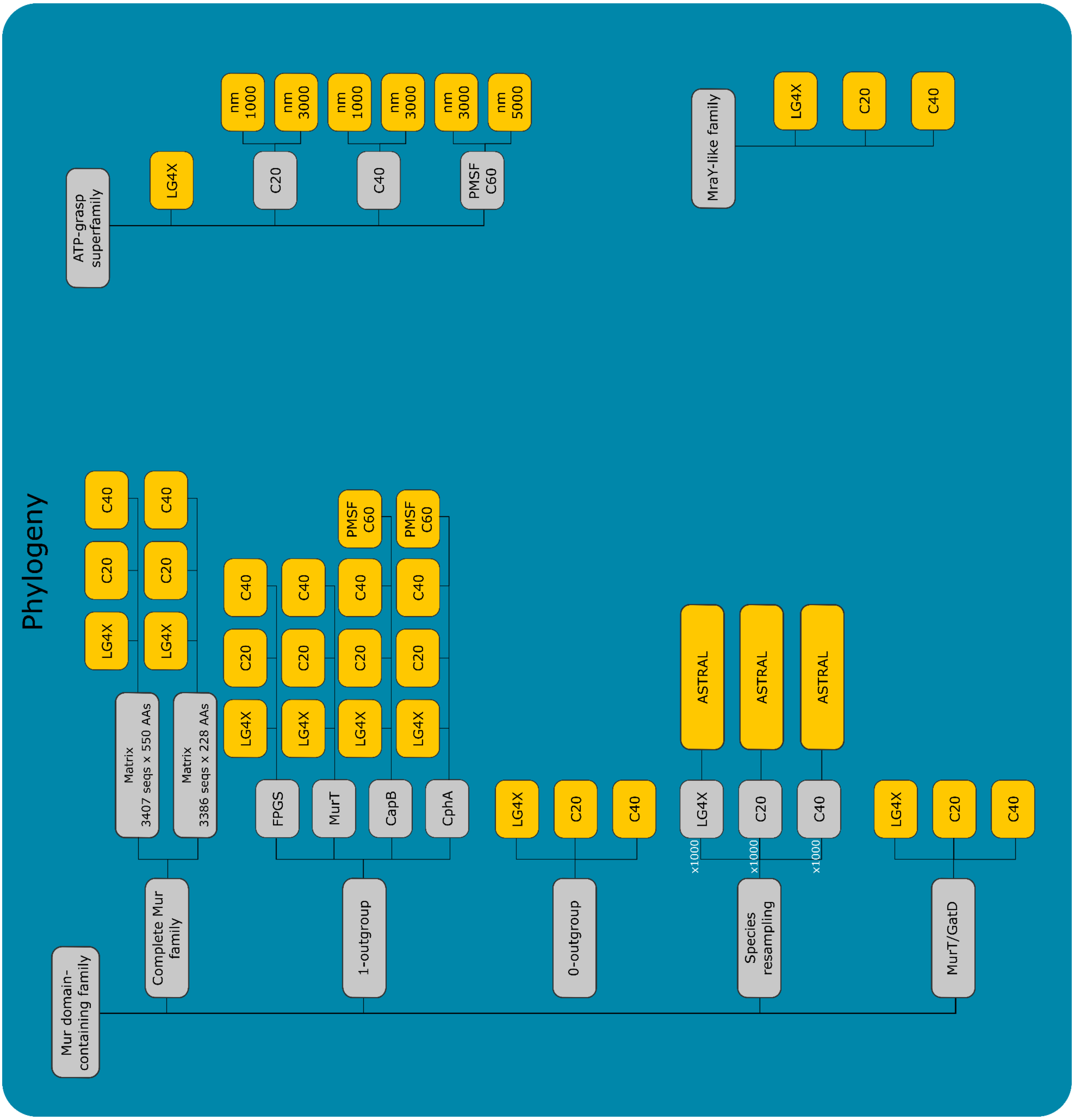

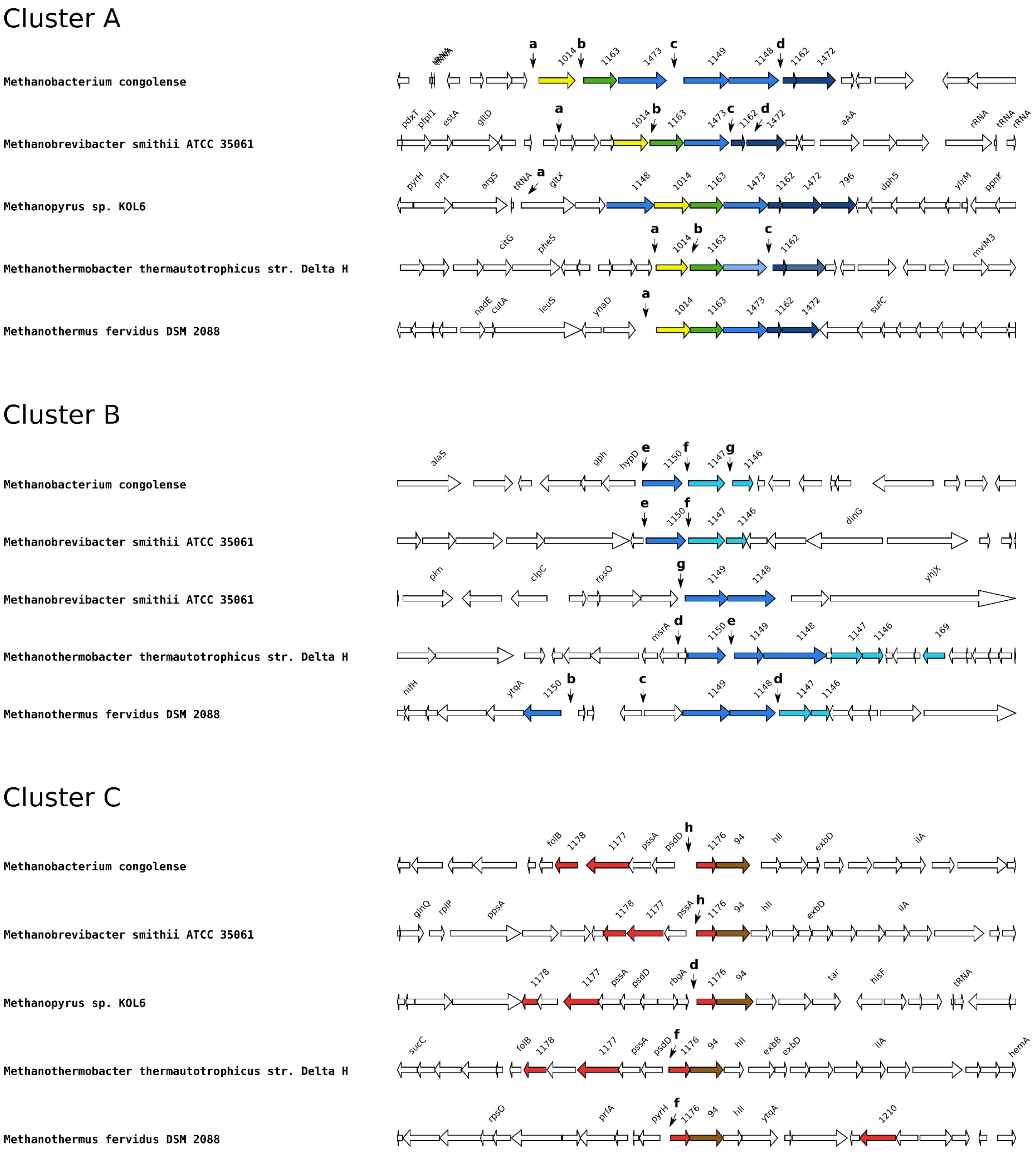
Cluster A
Methanobacterium congolense
tRNA
a
1014
b
1163
1473
c
1149
1148
d
1162
1472
Methanobrevibacter smithii ATCC 35061
pdxT
pfpI1
estA
gltD
a
1014
b
1163
1473
c
1162
d
1472
aAA
rRNA
tRNA
rRNA
Methanopyrus sp. KOL6
pyrH
prf1
argS
tRNA
a
gltX
1148
1014
1163
1473
1162
1472
796
dph5
ylxM
ppnK
Methanothermobacter thermautotrophicus str. Delta H
citG
pheS
a
1014
b
1163
c
1162
mviM3
Methanothermus fervidus DSM 2088
nadE
cutA
leuS
ynaD
a
1014
1163
1473
1162
1472
sufC
Cluster B
Methanobacterium congolense
alaS
gph
hypD
e
1150
f
1147
g
1146
Methanobrevibacter smithii ATCC 35061
e
1150
f
1147
1146
dinG
Methanobrevibacter smithii ATCC 35061
pkn
clpC
rpsO
g
1149
1148
yhjX
Methanothermobacter thermautotrophicus str. Delta H
msrA
d
1150
e
1149
1148
1147
1146
169
Methanothermus fervidus DSM 2088
nifH
ytqA
1150
b
c
1149
1148
d
1147
1146
Cluster C
Methanobacterium congolense
folB
1178
1177
pssA
psdD
h
1176
94
hll
exbD
ilA
Methanobrevibacter smithii ATCC 35061
glnQ
rplP
ppsA
1178
1177
pssA
h
1176
94
hll
exbD
ilA
Methanopyrus sp. KOL6
1178
1177
pssA
psdD
rbgA
d
1176
94
tar
hisF
tRNA
Methanothermobacter thermautotrophicus str. Delta H
sucC
folB
1178
1177
pssA
psdD
f
1176
94
hll
exbB
exbD
ilA
hemA
Methanothermus fervidus DSM 2088
rpsO
prfA
pyrH
f
1176
94
hll
ytqA
1210

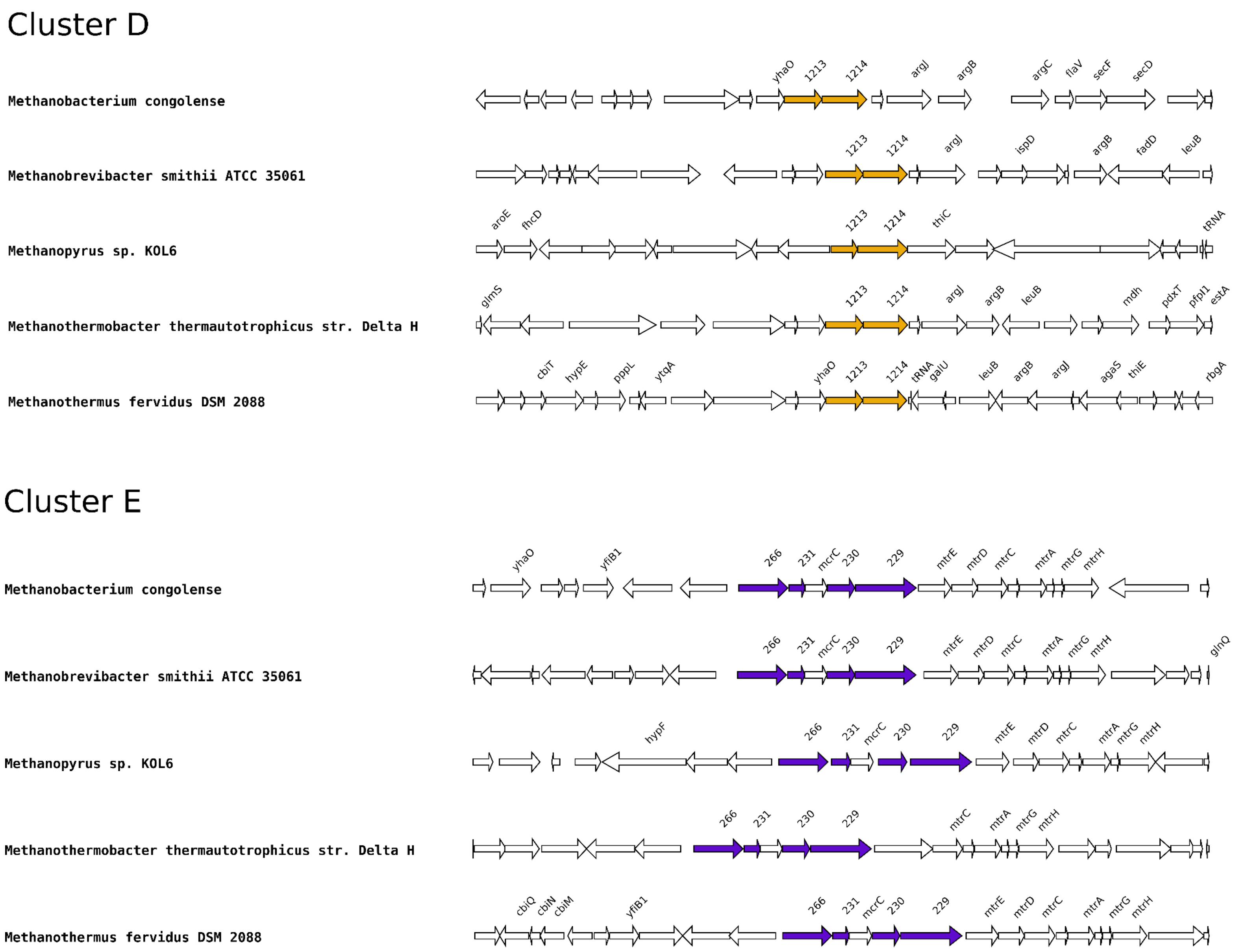


**Figure S2. Genetic organization of the five clusters.** Identified clustered genes are colored, whereas other genes are left in white. Above each identified gene is indicated its corresponding orthologous group (OG) number (without 'OG' and the following 0s). Intergenic regions used during the regulon pipeline (see Supplemental data) are indicated with a lowercase letter (Table S4). The intergenic regions "b" and "c" of *Methanopyrus* sp. are not shown in this figure and correspond to the upstream region of the OG0001150 and OG0001147, genes respectively.

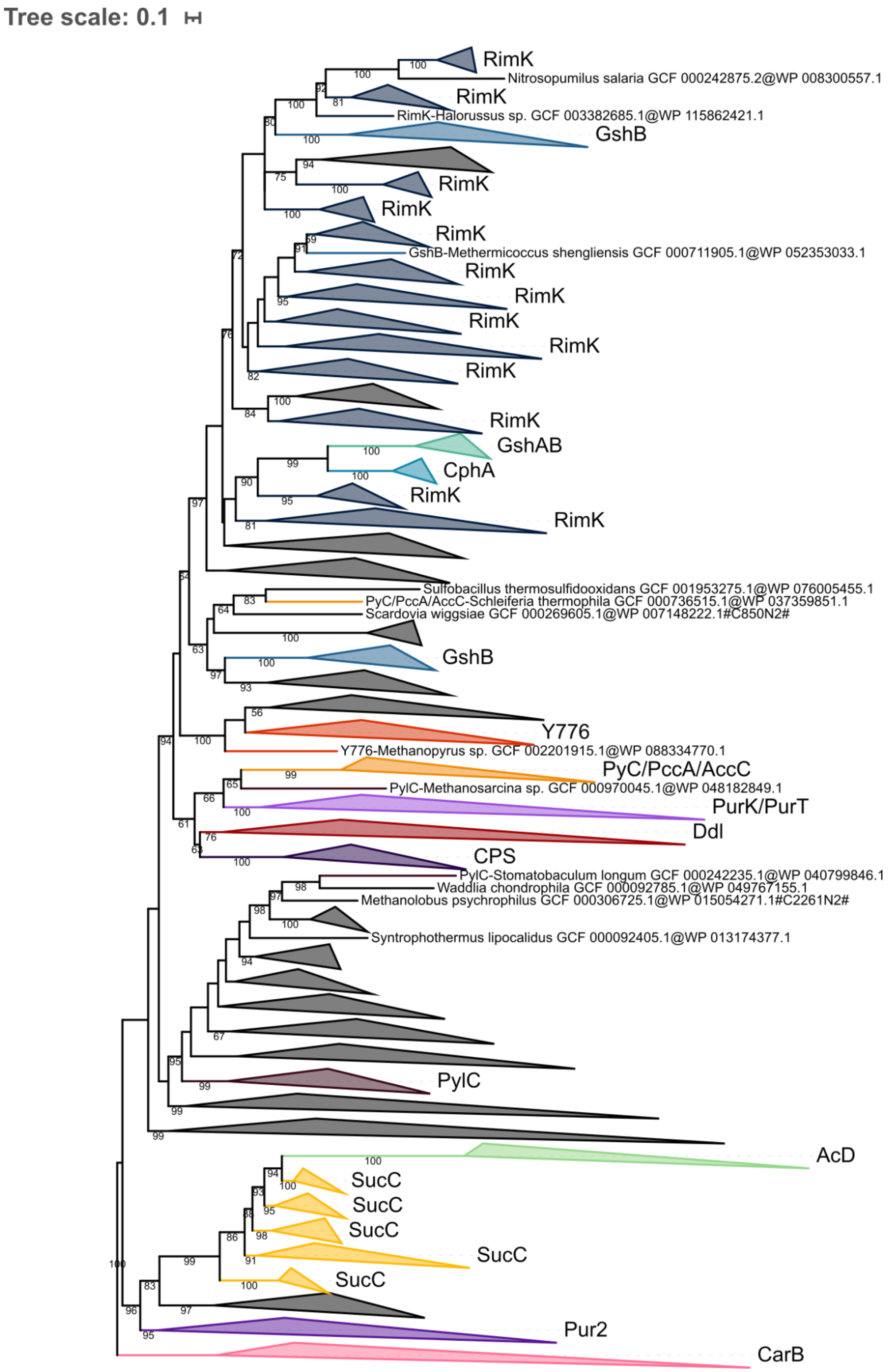

Tree scale: 0.1
RimK
Nitrosopumilus salaria GCF 000242875.2@WP 008300557.1
RimK
RimK-Halorussus sp. GCF 003382685.1@WP 115862421.1
GshB
RimK
RimK
RimK
GshB-Methermicoccus shengliensis GCF 000711905.1@WP 052353033.1
RimK
RimK
RimK
RimK
RimK
RimK
GshAB
CphA
RimK
RimK
Sulfobacillus thermosulfidooxidans GCF 001953275.1@WP 076005455.1
PyC/PccA/AccC-Schleiferia thermophila GCF 000736515.1@WP 037359851.1
Scardovia wiggsiae GCF 000269605.1@WP 007148222.1#C850N2#
GshB
Y776
Y776-Methanopyrus sp. GCF 002201915.1@WP 088334770.1
PyC/PccA/AccC
PylC-Methanosarcina sp. GCF 000970045.1@WP 048182849.1
PurK/PurT
Ddl
CPS
PylC-Stomatobaculum longum GCF 000242235.1@WP 040799846.1
Waddlia chondrophila GCF 000092785.1@WP 049767155.1
Methanolobus psychrophilus GCF 000306725.1@WP 015054271.1#C2261N2#
Syntrophothermus lipocalidus GCF 000092405.1@WP 013174377.1
PylC
AcD
SucC
SucC
SucC
SucC
SucC
Pur2
CarB

**Figure S3. Phylogenetic tree of the ATP-grasp superfamily rooted on CarB.** The tree was inferred from a matrix of 2,194 sequences x 180 unambiguously aligned AAs using IQ-TREE **under the C20+G4 model with 1000 iterations**. Tree visualization was performed using iTOL. Bootstrap support values are shown if greater or equal to 50. Branches were collapsed on homogeneous sequence annotation based on reference sequences. Here, OG0001014 is referred to as CPS. Black collapsed branches correspond to unannotated sequences.

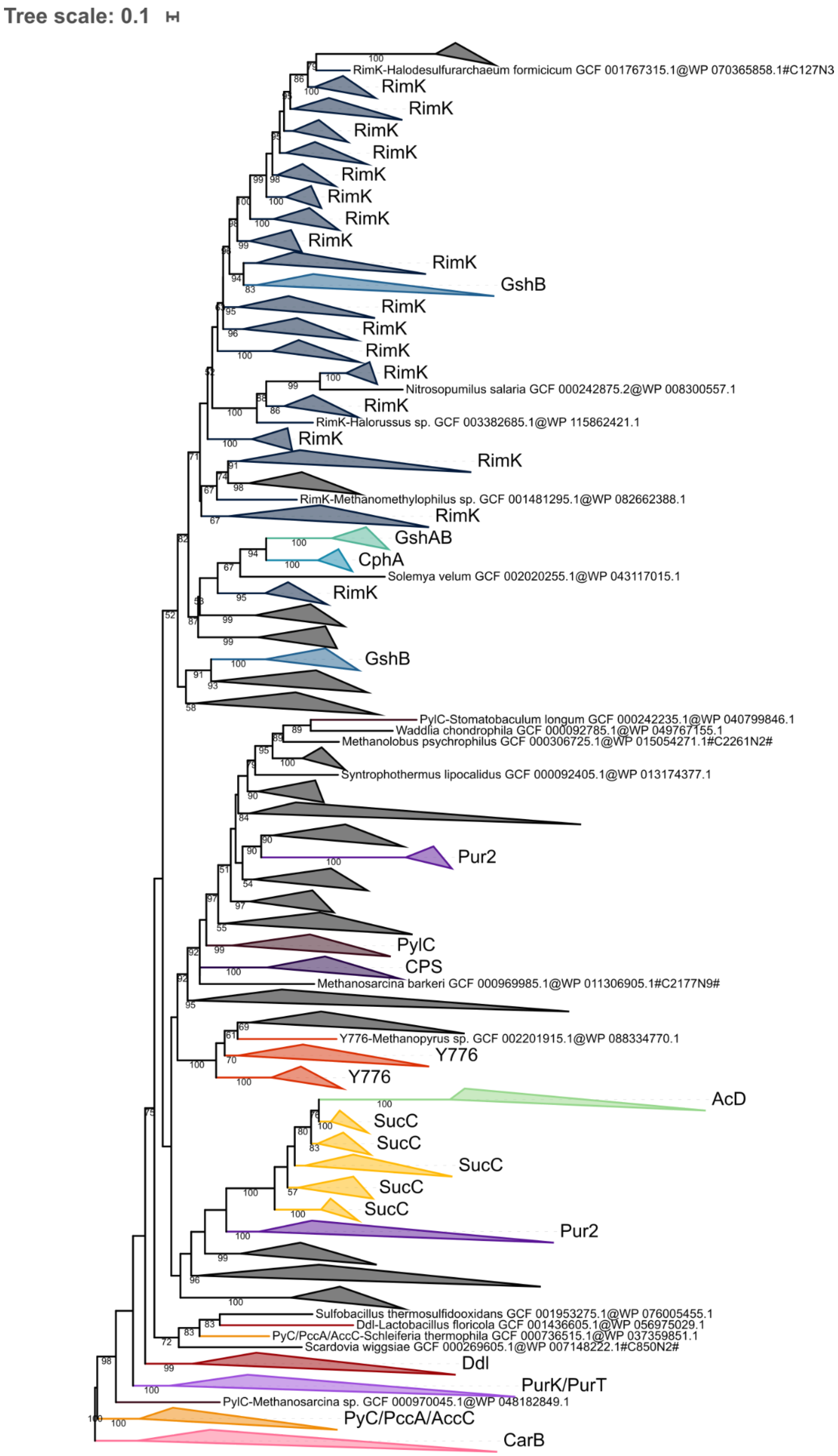
Tree scale: 0.1
RimK-Halodesulfurarchaeum formicicum GCF 001767315.1@WP 070365858.1#C127N3
RimK
RimK
RimK
RimK
RimK
RimK
RimK
RimK
RimK
GshB
RimK
RimK
RimK
RimK
Nitrosopumilus salaria GCF 000242875.2@WP 008300557.1
RimK
RimK-Halorussus sp. GCF 003382685.1@WP 115862421.1
RimK
RimK
RimK-Methanomethylophilus sp. GCF 001481295.1@WP 082662388.1
RimK
GshAB
CphA
Solemya velum GCF 002020255.1@WP 043117015.1
RimK
GshB
PylC-Stomatobaculum longum GCF 000242235.1@WP 040799846.1
Waddlia chondrophila GCF 000092785.1@WP 049767155.1
Methanolobus psychrophilus GCF 000306725.1@WP 015054271.1#C2261N2#
Syntrophothermus lipocalidus GCF 000092405.1@WP 013174377.1
Pur2
PylC
CPS
Methanosarcina barkeri GCF 000969985.1@WP 011306905.1#C2177N9#
Y776-Methanopyrus sp. GCF 002201915.1@WP 088334770.1
Y776
Y776
AcD
SucC
SucC
SucC
SucC
SucC
Pur2
Sulfobacillus thermosulfidooxidans GCF 001953275.1@WP 076005455.1
Ddl-Lactobacillus floricola GCF 001436605.1@WP 056975029.1
PyC/PccA/AccC-Schleiferia thermophila GCF 000736515.1@WP 037359851.1
Scardovia wiggsiae GCF 000269605.1@WP 007148222.1#C850N2#
Ddl
PurK/PurT
PylC-Methanosarcina sp. GCF 000970045.1@WP 048182849.1
PyC/PccA/AccC
CarB

**Figure S4. Phylogenetic tree of the ATP-grasp superfamily rooted on CarB.** The tree was inferred from a matrix of 2,194 sequences x 180 unambiguously aligned AAs using IQ-TREE **under the C20+G4 model with 3000 iterations**. Tree visualization was performed using iTOL. Bootstrap support values are shown if greater or equal to 50. Branches were collapsed on homogeneous sequence annotation based on reference sequences. Here, OG0001014 is referred to as CPS. Black collapsed branches correspond to unannotated sequences.

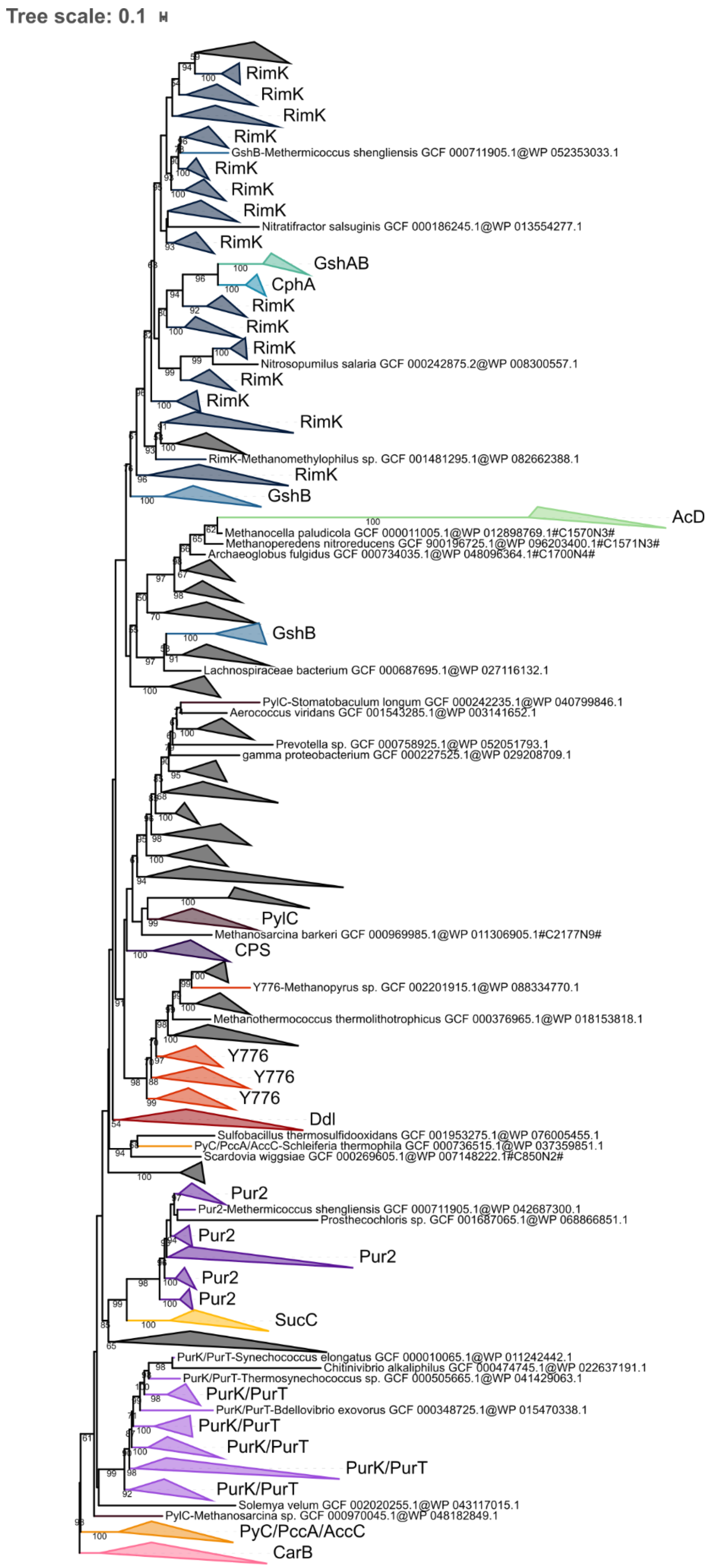
Tree scale: 0.1
RimK
RimK
RimK
RimK
GshB-Methermicoccus shengliensis GCF 000711905.1@WP 052353033.1
RimK
RimK
RimK
Nitratifractor salsuginis GCF 000186245.1@WP 013554277.1
RimK
GshAB
CphA
RimK
RimK
RimK
Nitrosopumilus salaria GCF 000242875.2@WP 008300557.1
RimK
RimK
RimK
RimK-Methanomethylophilus sp. GCF 001481295.1@WP 082662388.1
RimK
GshB
AcD
Methanocella paludicola GCF 000011005.1@WP 012898769.1#C1570N3#
Methanoperedens nitroreducens GCF 900196725.1@WP 096203400.1#C1571N3#
Archaeoglobus fulgidus GCF 000734035.1@WP 048096364.1#C1700N4#
GshB
Lachnospiraceae bacterium GCF 000687695.1@WP 027116132.1
PylC-Stomatobaculum longum GCF 000242235.1@WP 040799846.1
Aerococcus viridans GCF 001543285.1@WP 003141652.1
Prevotella sp. GCF 000758925.1@WP 052051793.1
gamma proteobacterium GCF 000227525.1@WP 029208709.1
PylC
Methanosarcina barkeri GCF 000969985.1@WP 011306905.1#C2177N9#
CPS
Y776-Methanopyrus sp. GCF 002201915.1@WP 088334770.1
Methanothermococcus thermolithotrophicus GCF 000376965.1@WP 018153818.1
Y776
Y776
Y776
Ddl
Sulfobacillus thermosulfidooxidans GCF 001953275.1@WP 076005455.1
PyC/PccA/AccC-Schleiferia thermophila GCF 000736515.1@WP 037359851.1
Scardovia wiggsiae GCF 000269605.1@WP 007148222.1#C850N2#
Pur2
Pur2-Methermicoccus shengliensis GCF 000711905.1@WP 042687300.1
Prosthecochloris sp. GCF 001687065.1@WP 068866851.1
Pur2
Pur2
Pur2
Pur2
SucC
PurK/PurT-Synechococcus elongatus GCF 000010065.1@WP 011242442.1
Chitinivibrio alkaliphilus GCF 000474745.1@WP 022637191.1
PurK/PurT-Thermosynechococcus sp. GCF 000505665.1@WP 041429063.1
PurK/PurT
PurK/PurT-Bdellovibrio exovorus GCF 000348725.1@WP 015470338.1
PurK/PurT
PurK/PurT
PurK/PurT
PurK/PurT
Solemya velum GCF 002020255.1@WP 043117015.1
PylC-Methanosarcina sp. GCF 000970045.1@WP 048182849.1
PyC/PccA/AccC
CarB

**Figure S5. Phylogenetic tree of the ATP-grasp superfamily rooted on CarB.** The tree was inferred from a matrix of 2,194 sequences x 180 unambiguously aligned AAs using IQ-TREE **under the C40+G4 model with 1000 iterations**. Tree visualization was performed using iTOL. Bootstrap support values are shown if greater or equal to 50. Branches were collapsed on homogeneous sequence annotation based on reference sequences. Here, OG0001014 is referred to as CPS. Black collapsed branches correspond to unannotated sequences.

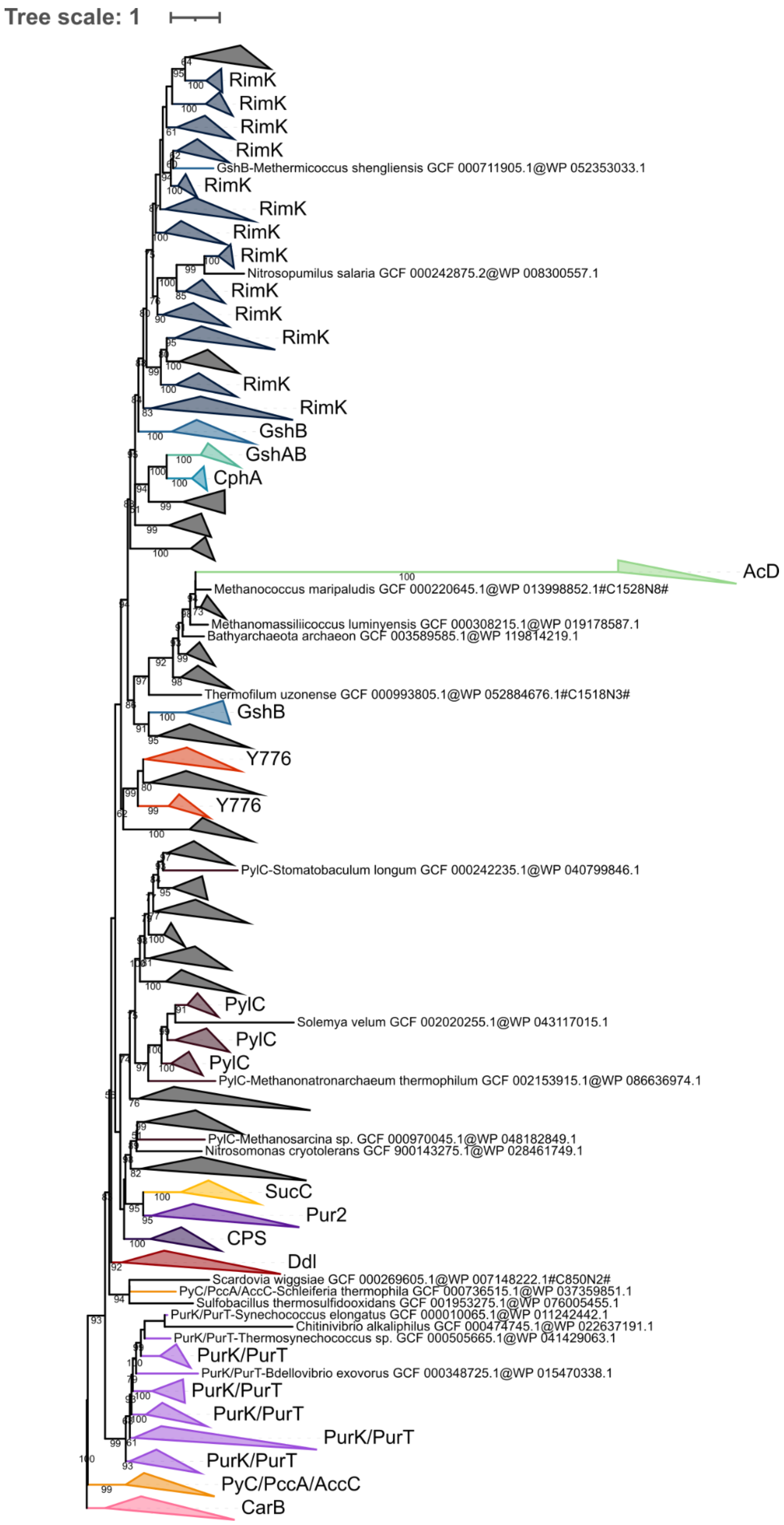
Tree scale: 1
RimK
RimK
RimK
RimK
GshB-Methermicoccus shengliensis GCF 000711905.1@WP 052353033.1
RimK
RimK
RimK
RimK
Nitrosopumilus salaria GCF 000242875.2@WP 008300557.1
RimK
RimK
RimK
RimK
RimK
GshB
GshAB
CphA
AcD
Methanococcus maripaludis GCF 000220645.1@WP 013998852.1#C1528N8#
Methanomassiliicoccus luminyensis GCF 000308215.1@WP 019178587.1
Bathyarchaeota archaeon GCF 003589585.1@WP 119814219.1
Thermofilum uzonense GCF 000993805.1@WP 052884676.1#C1518N3#
GshB
Y776
Y776
PylC-Stomatobaculum longum GCF 000242235.1@WP 040799846.1
PylC
Solemya velum GCF 002020255.1@WP 043117015.1
PylC
PylC
PylC-Methanonatronarchaeum thermophilum GCF 002153915.1@WP 086636974.1
PylC-Methanosarcina sp. GCF 000970045.1@WP 048182849.1
Nitrosomonas cryotolerans GCF 900143275.1@WP 028461749.1
SucC
Pur2
CPS
Ddl
Scardovia wiggsiae GCF 000269605.1@WP 007148222.1#C850N2#
PyC/PccA/AccC-Schleiferia thermophila GCF 000736515.1@WP 037359851.1
Sulfobacillus thermosulfidooxidans GCF 001953275.1@WP 076005455.1
PurK/PurT-Synechococcus elongatus GCF 000010065.1@WP 011242442.1
Chitinivibrio alkaliphilus GCF 000474745.1@WP 022637191.1
PurK/PurT-Thermosynechococcus sp. GCF 000505665.1@WP 041429063.1
PurK/PurT
PurK/PurT-Bdellovibrio exovorus GCF 000348725.1@WP 015470338.1
PurK/PurT
PurK/PurT
PurK/PurT
PurK/PurT
PyC/PccA/AccC
CarB

**Figure S6. Phylogenetic tree of the ATP-grasp superfamily rooted on CarB.** The tree was inferred from a matrix of 2,194 sequences x 180 unambiguously aligned AAs using IQ-TREE **under the C40+G4 model with 3000 iterations**. Tree visualization was performed using iTOL. Bootstrap support values are shown if greater or equal to 50. Branches were collapsed on homogeneous sequence annotation based on reference sequences. Here, OG0001014 is referred to as CPS. Black collapsed branches correspond to unannotated sequences.

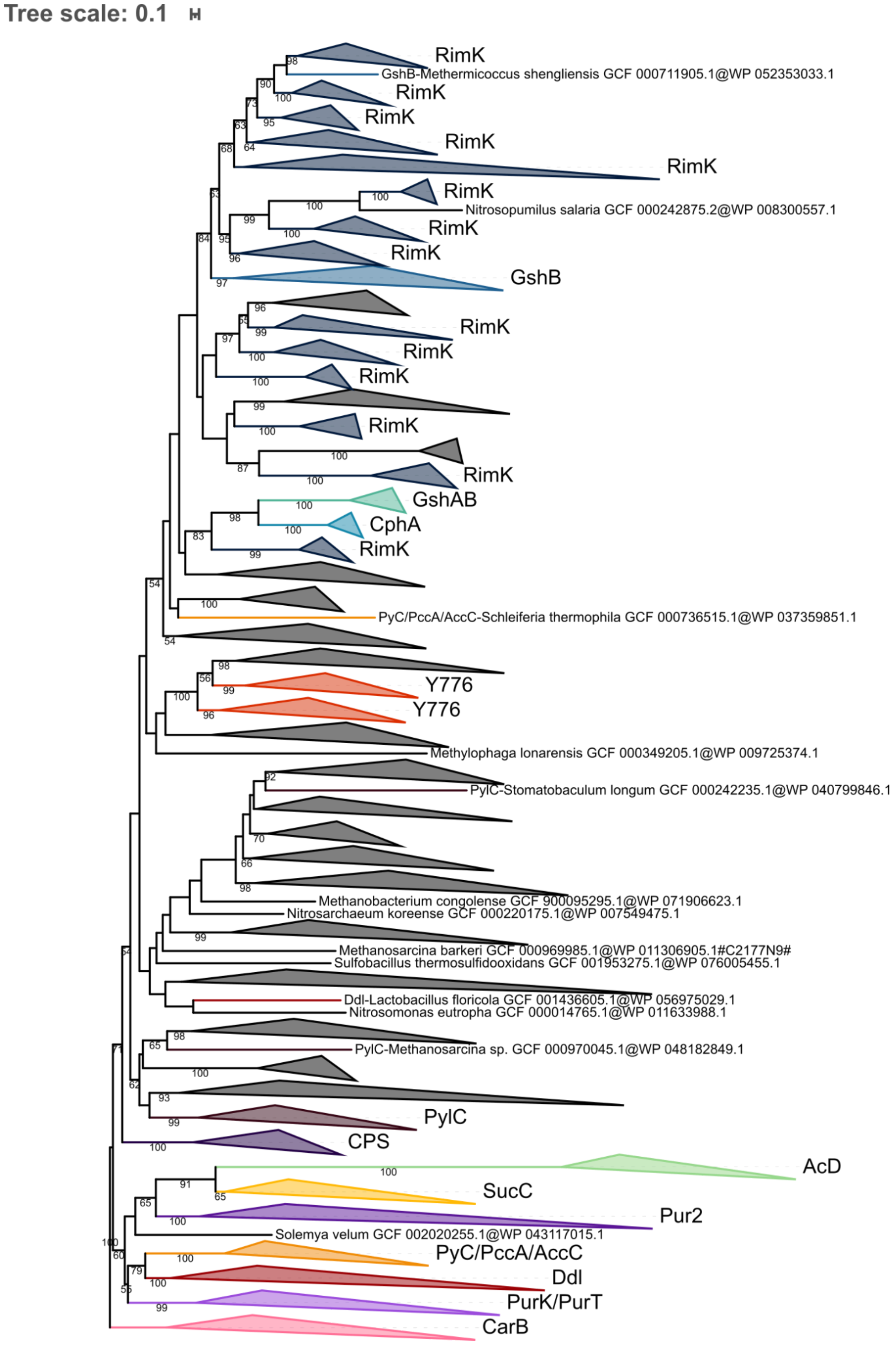
Tree scale: 0.1
RimK
GshB-Methermicoccus shengliensis GCF 000711905.1@WP 052353033.1
RimK
RimK
RimK
RimK
RimK
Nitrosopumilus salaria GCF 000242875.2@WP 008300557.1
RimK
RimK
GshB
RimK
RimK
RimK
RimK
RimK
GshAB
CphA
RimK
PyC/PccA/AccC-Schleiferia thermophila GCF 000736515.1@WP 037359851.1
Y776
Y776
Methylophaga lonarensis GCF 000349205.1@WP 009725374.1
PylC-Stomatobaculum longum GCF 000242235.1@WP 040799846.1
Methanobacterium congolense GCF 900095295.1@WP 071906623.1
Nitrosarchaeum koreense GCF 000220175.1@WP 007549475.1
Methanosarcina barkeri GCF 000969985.1@WP 011306905.1#C2177N9#
Sulfobacillus thermosulfidooxidans GCF 001953275.1@WP 076005455.1
Ddl-Lactobacillus floricola GCF 001436605.1@WP 056975029.1
Nitrosomonas eutropha GCF 000014765.1@WP 011633988.1
PylC-Methanosarcina sp. GCF 000970045.1@WP 048182849.1
PylC
CPS
AcD
SucC
Pur2
Solemya velum GCF 002020255.1@WP 043117015.1
PyC/PccA/AccC
Ddl
PurK/PurT
CarB

**Figure S7. Phylogenetic tree of the ATP-grasp superfamily rooted on CarB.** The tree was inferred from a matrix of 2,194 sequences x 180 unambiguously aligned AAs using IQ-TREE **under the PMSF LG+C60+G4 model with 3000 iterations**. Tree visualization was performed using iTOL. Bootstrap support values are shown if greater or equal to 50. Branches were collapsed on homogeneous sequence annotation based on reference sequences. Here, OG0001014 is referred to as CPS. Black collapsed branches correspond to unannotated sequences.

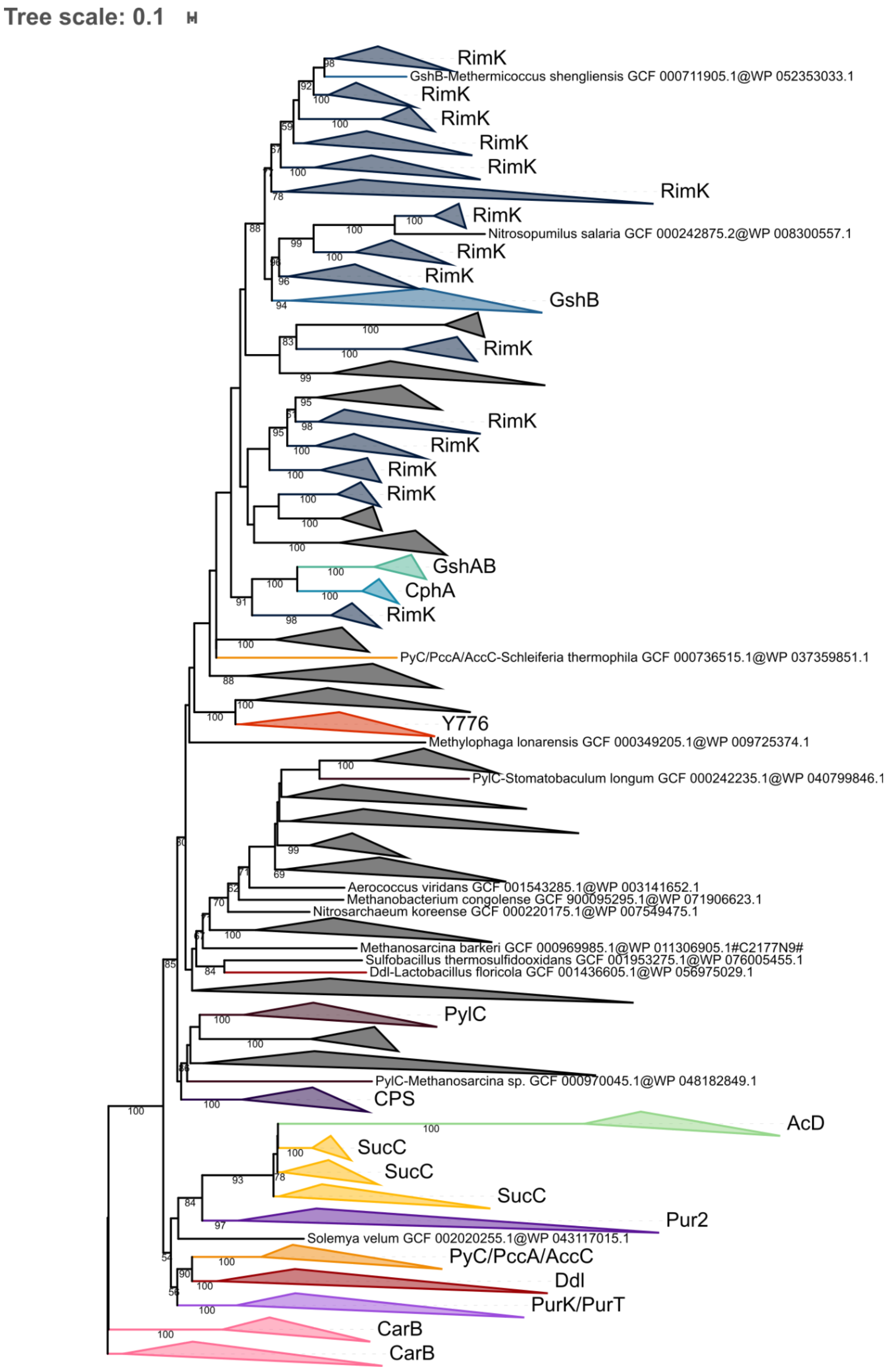
Tree scale: 0.1
RimK
GshB-Methermicoccus shengliensis GCF 000711905.1@WP 052353033.1
RimK
RimK
RimK
RimK
RimK
RimK
Nitrosopumilus salaria GCF 000242875.2@WP 008300557.1
RimK
RimK
GshB
RimK
RimK
RimK
RimK
RimK
GshAB
CphA
RimK
PyC/PccA/AccC-Schleiferia thermophila GCF 000736515.1@WP 037359851.1
Y776
Methylophaga lonarensis GCF 000349205.1@WP 009725374.1
PylC-Stomatobaculum longum GCF 000242235.1@WP 040799846.1
Aerococcus viridans GCF 001543285.1@WP 003141652.1
Methanobacterium congolense GCF 900095295.1@WP 071906623.1
Nitrosarchaeum koreense GCF 000220175.1@WP 007549475.1
Methanosarcina barkeri GCF 000969985.1@WP 011306905.1#C2177N9#
Sulfobacillus thermosulfidooxidans GCF 001953275.1@WP 076005455.1
Ddl-Lactobacillus floricola GCF 001436605.1@WP 056975029.1
PylC
PylC-Methanosarcina sp. GCF 000970045.1@WP 048182849.1
CPS
AcD
SucC
SucC
SucC
Pur2
Solemya velum GCF 002020255.1@WP 043117015.1
PyC/PccA/AccC
Ddl
PurK/PurT
CarB
CarB

**Figure S8. Phylogenetic tree of the ATP-grasp superfamily rooted on CarB.** The tree was inferred from a matrix of 2,194 sequences x 180 unambiguously aligned AAs using IQ-TREE **under the PMSF LG+C60+G4 model with 5000 iterations**. Tree visualization was performed using iTOL. Bootstrap support values are shown if greater or equal to 50. Branches were collapsed on homogeneous sequence annotation based on reference sequences. Here, OG0001014 is referred to as CPS. Black collapsed branches correspond to unannotated sequences.

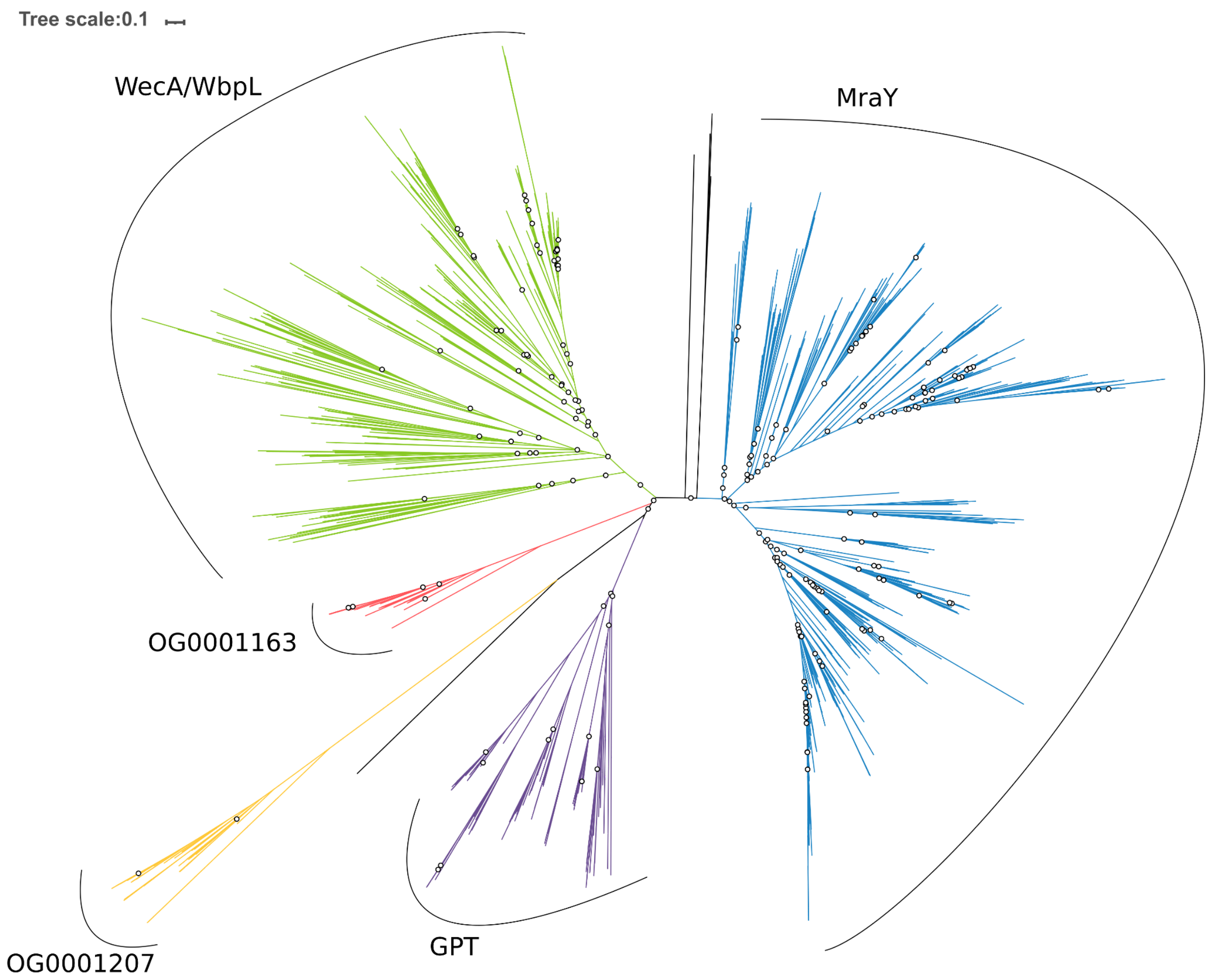


**Figure S9. Unrooted phylogenetic tree of the MraY-like family.** The tree was constructed from a matrix of 1,070 sequences x 408 unambiguously aligned AAs using IQ-TREE **under the LG4X+R4 model**. Open circles correspond to bootstrap support values lower than 90. Blue sequences correspond to a MraY annotation, green to WecA/WbpL, red to OG0001163 (MraY-like), yellow to OG0001207, purple to GPT, and black to six unannotated bacterial sequences. Regarding those unannotated sequences, the sequence of *Syntrophaceticus schinkii* is always basal to OG0001207, whereas the group composed of two sequences of

*Ruminococcaceae* sp. and two sequences of *Treponema* sp. is always basal to MraY. The last sequence from *Ruminococcus* sp. is basal to MraY.

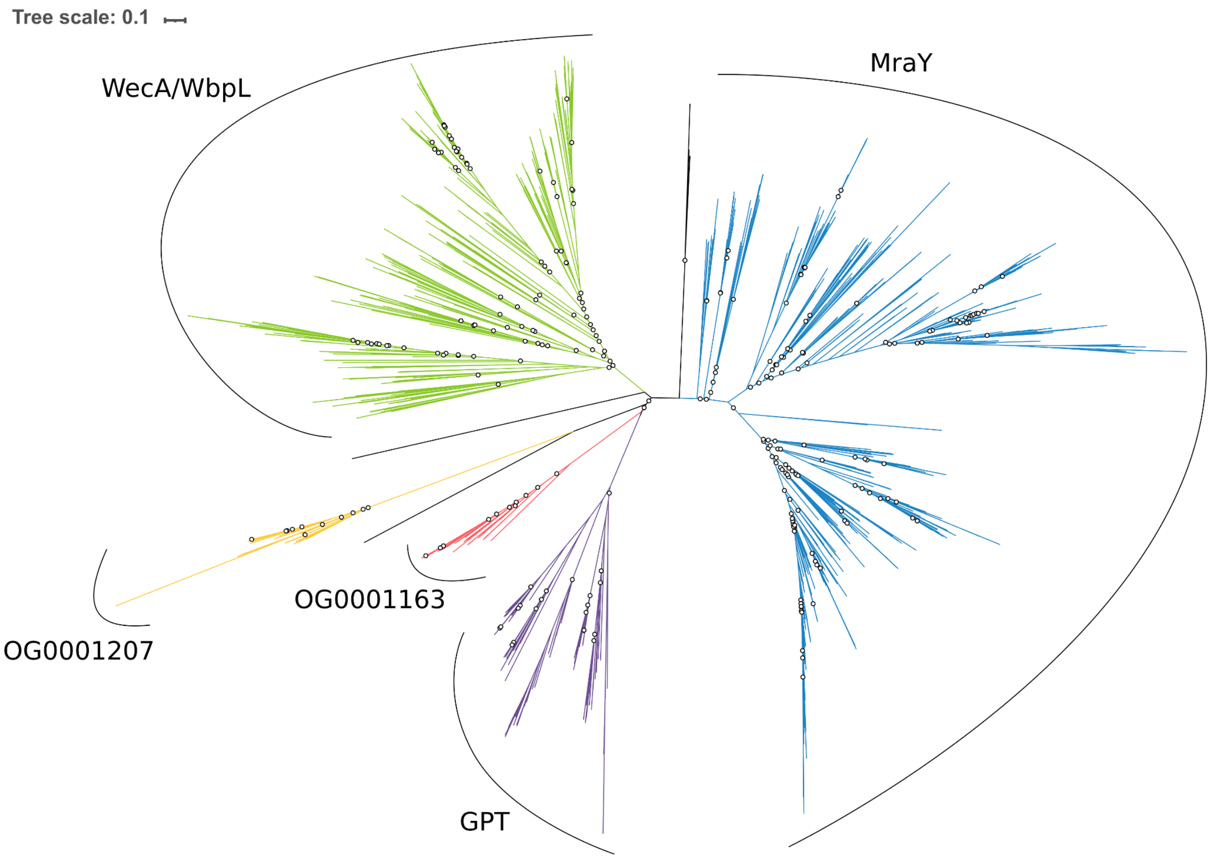


**Figure S10. Unrooted phylogenetic tree of the MraY-like family.** The tree was constructed from a matrix of 1,070 sequences x 408 unambiguously aligned AAs using IQ-TREE **under the C20+G4 model**. Open circles correspond to bootstrap support values lower than 90. Blue sequences correspond to a MraY annotation, green to WecA/WbpL, red to OG0001163 (MraY-like), yellow to OG0001207, purple to GPT, and black to six unannotated bacterial sequences. Regarding those unannotated sequences, the sequence of *Syntrophaceticus schinkii* is always basal to OG0001207, whereas the group composed of two sequences of *Ruminococcaceae* sp. and two sequences of *Treponema* sp. is always basal to MraY. The last sequence from *Ruminococcus* sp. is basal to WecA/WbpL.

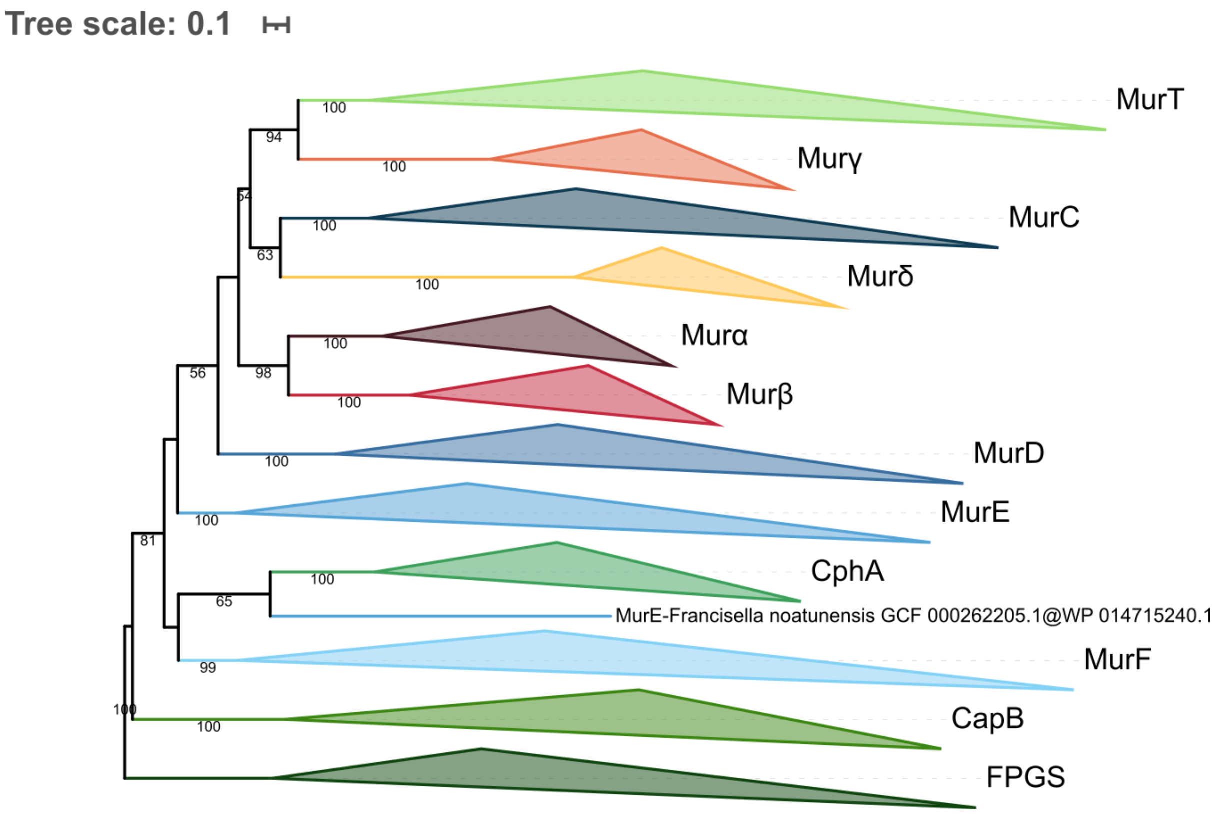


**Figure S11. Phylogenetic tree of the Mur domain-containing family rooted on FPGS.** The tree was inferred from a matrix of **3407 sequences x 550 unambiguously aligned AAs** using IQ-TREE **under the LG4X+R4 model**. Tree visualization was performed using iTOL. Bootstrap values are shown if greater or equal to 50. Branches were collapsed on homogeneous sequence annotation.

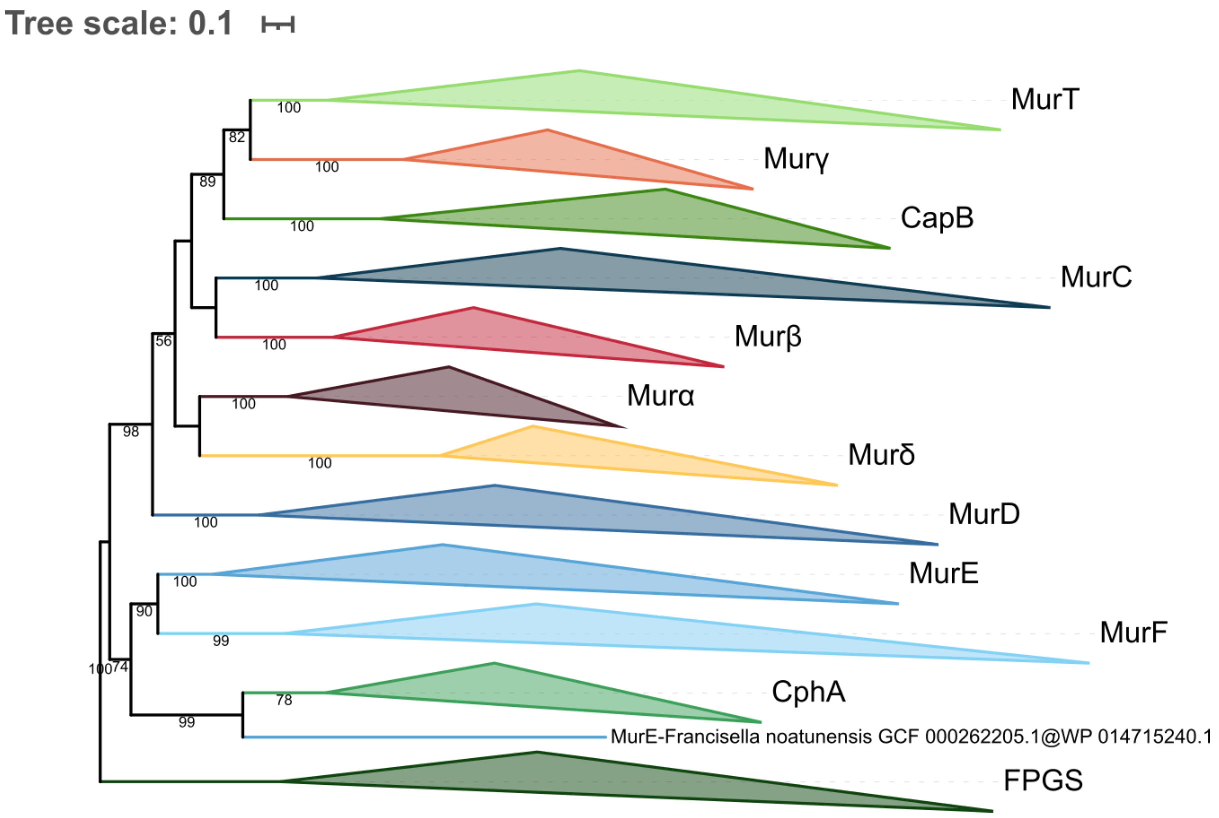


**Figure S12. Phylogenetic tree of the Mur domain-containing family rooted on FPGS.** The tree was inferred from a matrix of **3407 sequences x 550 unambiguously aligned AAs** using IQ-TREE **under the C20+G4 model**. Tree visualization was performed using iTOL. Bootstrap values are shown if greater or equal to 50. Branches were collapsed on homogeneous sequence annotation.

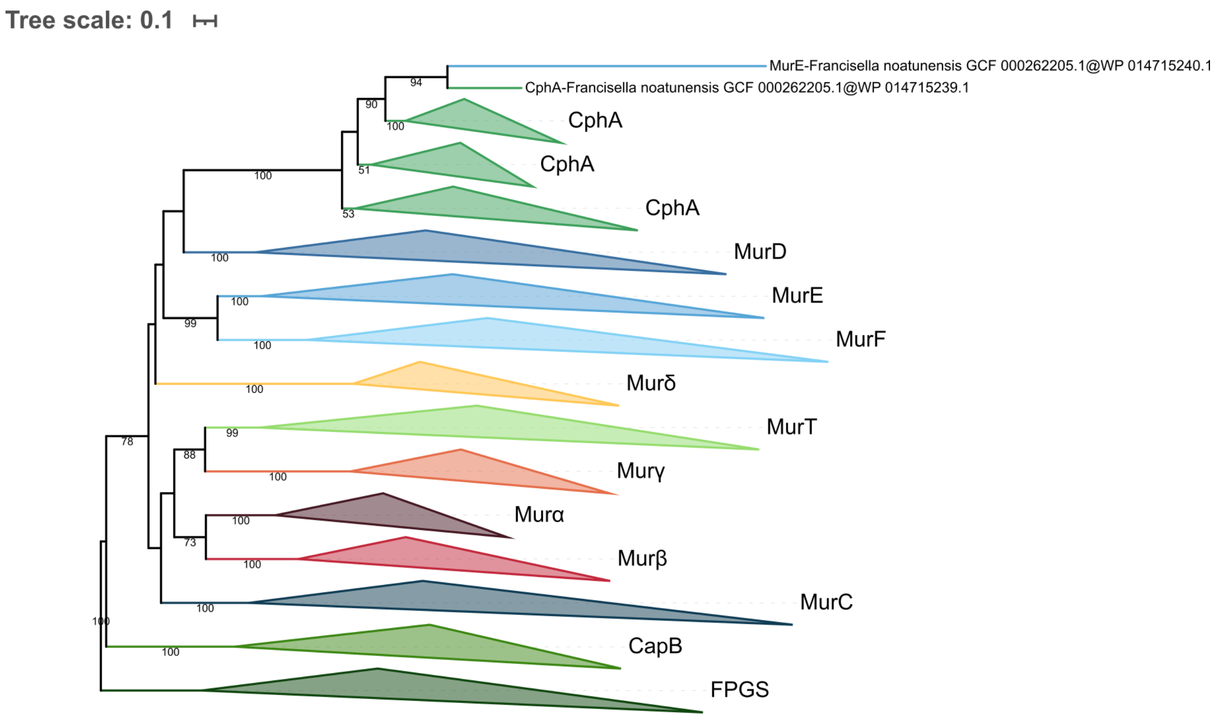


**Figure S13. Phylogenetic tree of the Mur domain-containing family rooted on FPGS.** The tree was inferred from a matrix of **3407 sequences x 550 unambiguously aligned AAs** using IQ-TREE **under the C40+G4 model**. Tree visualization was performed using iTOL. Bootstrap values are shown if greater or equal to 50. Branches were collapsed on homogeneous sequence annotation.

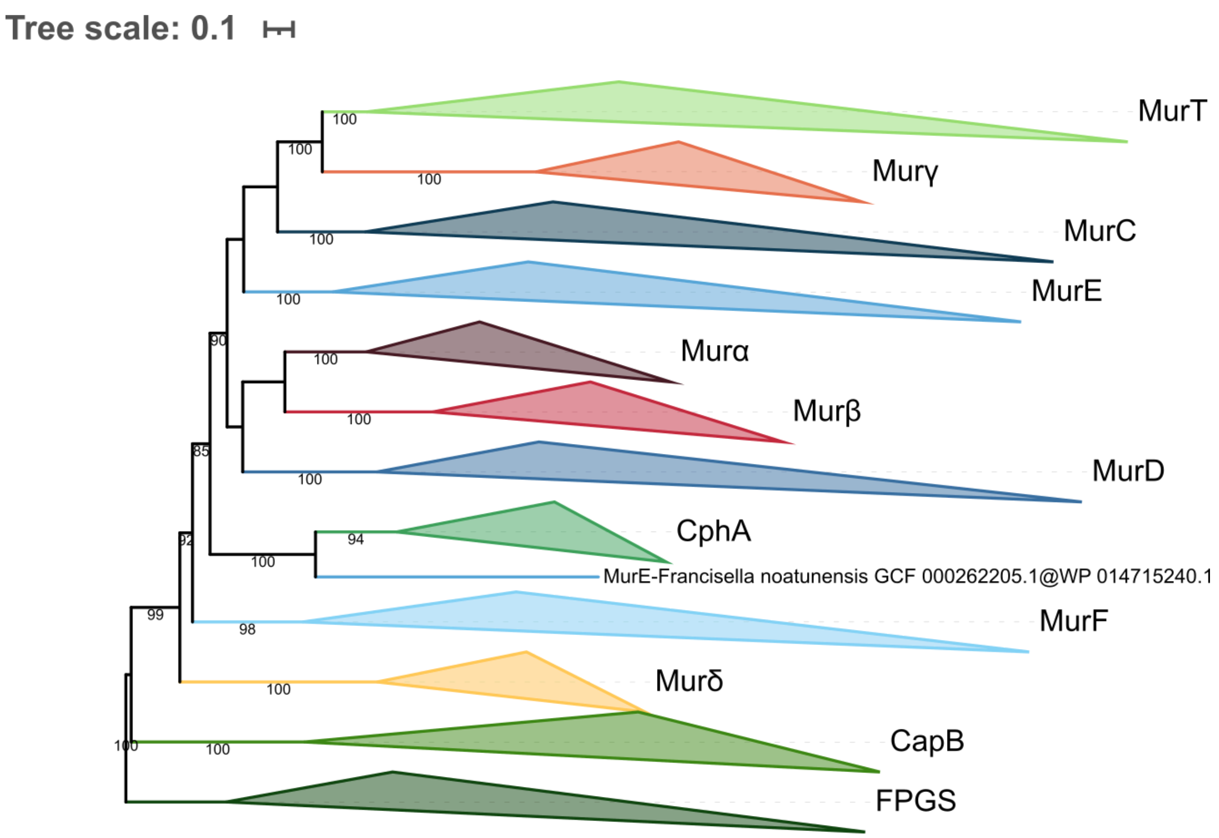


**Figure S14. Phylogenetic tree of the Mur domain-containing family rooted on FPGS.** The tree was inferred from a matrix of **3386 sequences x 228 unambiguously aligned AAs** using IQ-TREE **under the LG4X+R4 model**. Tree visualization was performed using iTOL. Bootstrap values are shown if greater or equal to 50. Branches were collapsed on homogeneous sequence annotation.

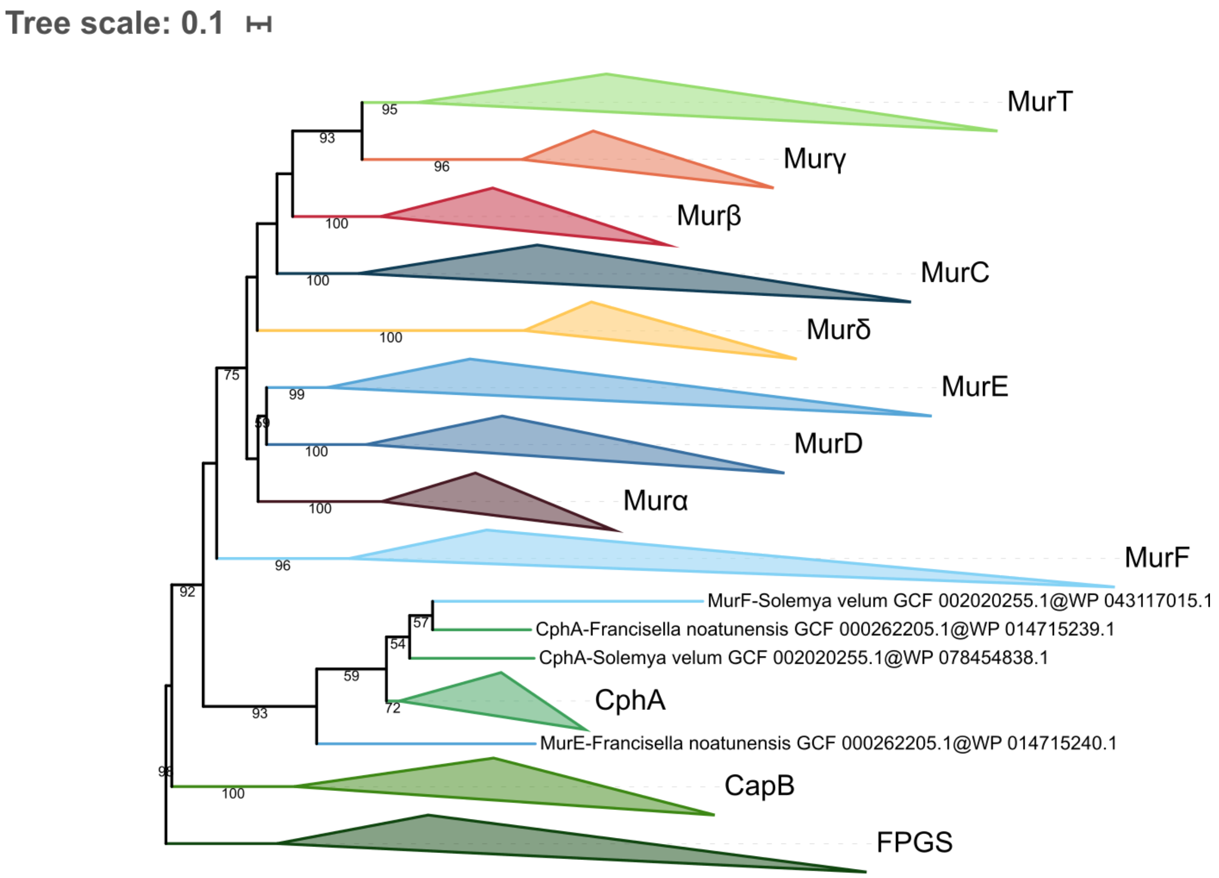


**Figure S15. Phylogenetic tree of the Mur domain-containing family rooted on FPGS.** The tree was inferred from a matrix of **3386 sequences x 228 unambiguously aligned AAs** using IQ-TREE **under the C20+G4 model**. Tree visualization was performed using iTOL. Bootstrap values are shown if greater or equal to 50. Branches were collapsed on homogeneous sequence annotation.

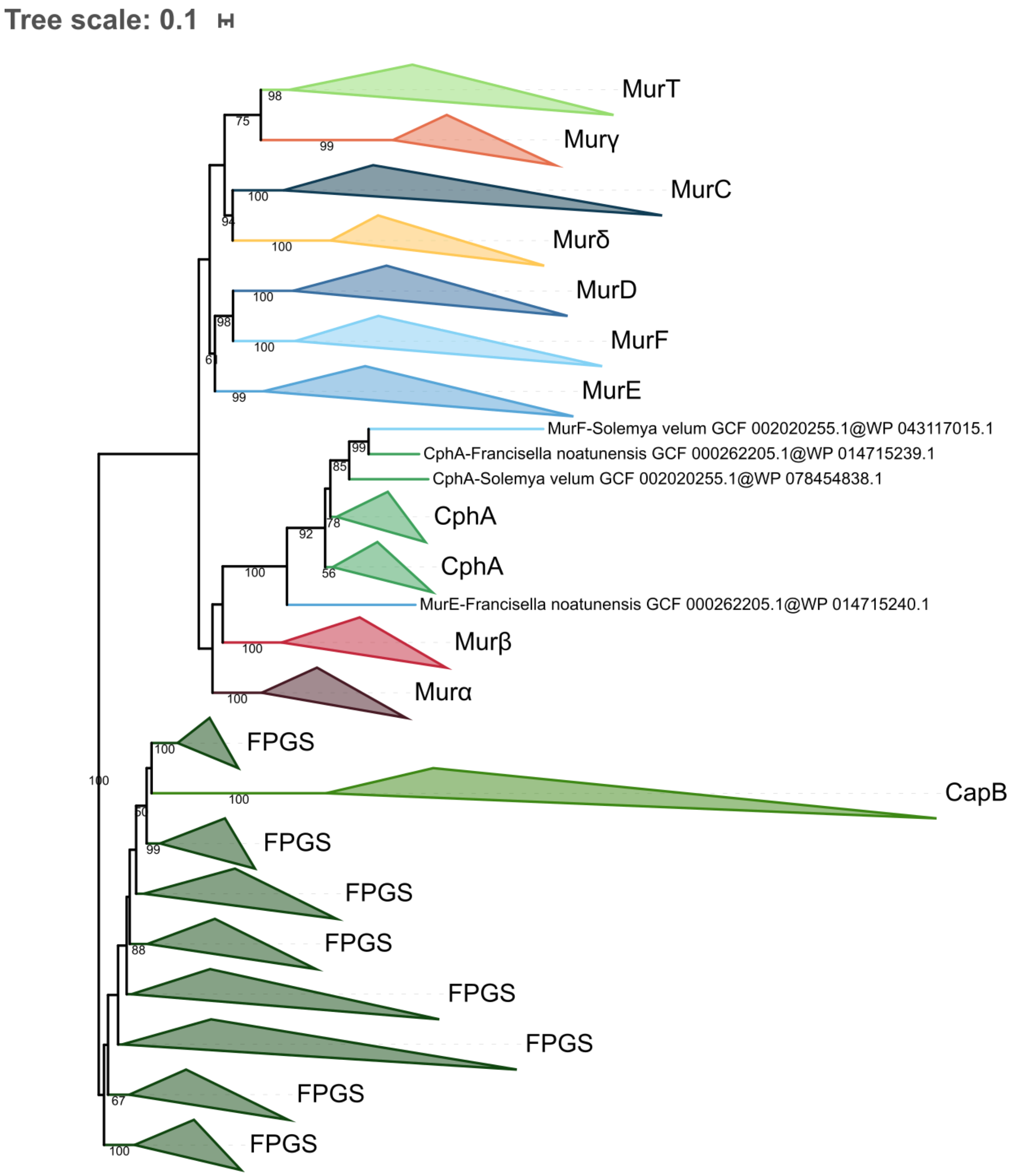


**Figure S16. Phylogenetic tree of the Mur domain-containing family rooted on FPGS.** The tree was inferred from a matrix of **3386 sequences x 228 unambiguously aligned AAs** using IQ-TREE **under the C40+G4 model**. Tree visualization was performed using iTOL. Bootstrap values are shown if greater or equal to 50. Branches were collapsed on homogeneous sequence annotation.

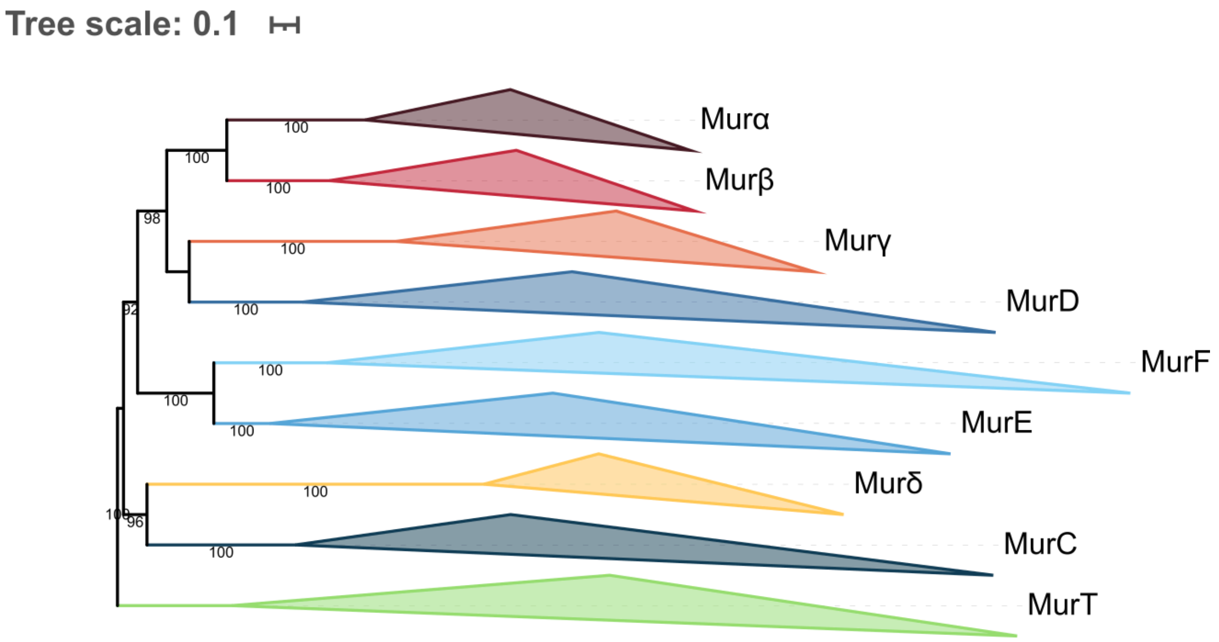


**Figure S17. Phylogenetic tree of the Mur domain-containing family rooted on MurT.** The tree was inferred from a matrix of **2677 sequences x 525 unambiguously aligned AAs** using IQ-TREE **under the LG4X+R4 model**. Tree visualization was performed using iTOL. Bootstrap values are shown if greater or equal to 50. Branches were collapsed on homogeneous sequence annotation.

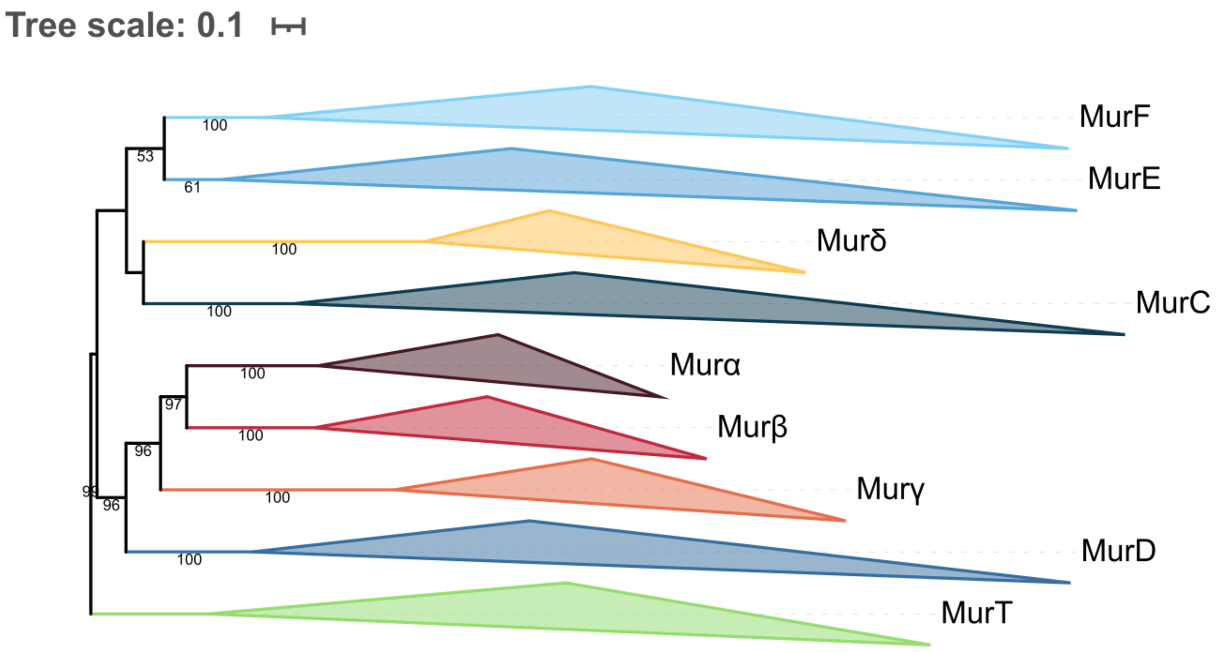


**Figure S18. Phylogenetic tree of the Mur domain-containing family rooted on MurT.** The tree was inferred from a matrix of **2677 sequences x 525 unambiguously aligned AAs** using IQ-TREE **under the C20+G4 model**. Tree visualization was performed using iTOL. Bootstrap values are shown if greater or equal to 50. Branches were collapsed on homogeneous sequence annotation.

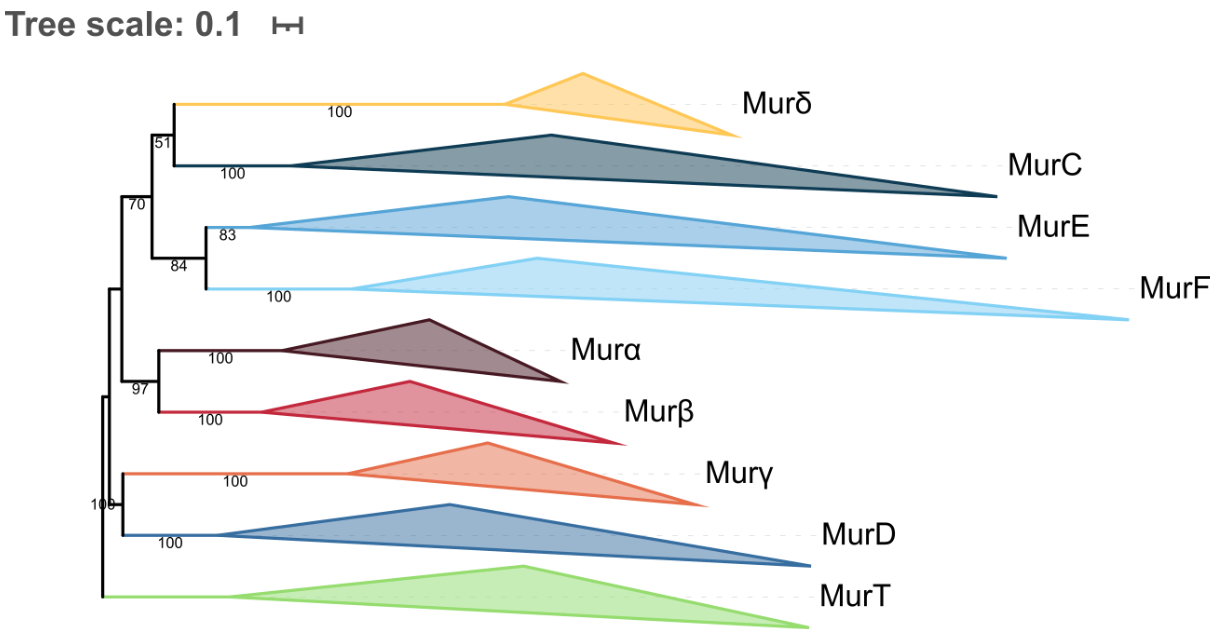


**Figure S19. Phylogenetic tree of the Mur domain-containing family rooted on MurT.** The tree was inferred from a matrix of **2677 sequences x 525 unambiguously aligned AAs** using IQ-TREE **under the C40+G4 model**. Tree visualization was performed using iTOL. Bootstrap values are shown if greater or equal to 50. Branches were collapsed on homogeneous sequence annotation.

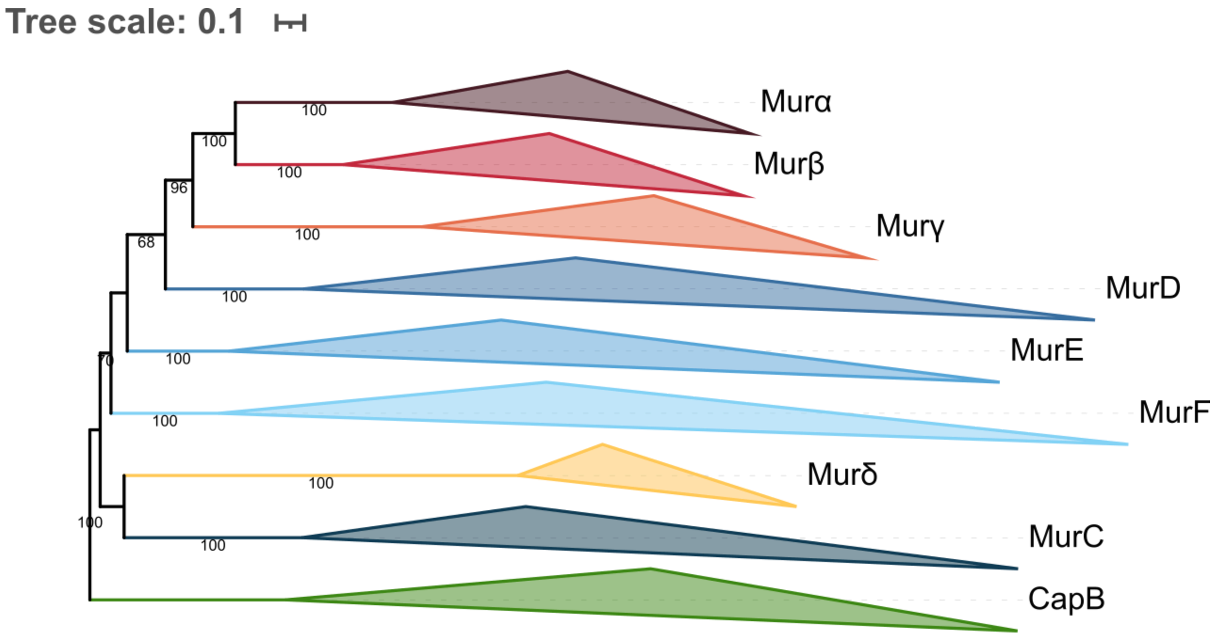


**Figure S20. Phylogenetic tree of the Mur domain-containing family rooted on CapB.** The tree was inferred from a matrix of **2519 sequences x 532 unambiguously aligned AAs** using IQ-TREE **under the LG4X+R4 model**. Tree visualization was performed using iTOL. Bootstrap values are shown if greater or equal to 50. Branches were collapsed on homogeneous sequence annotation.

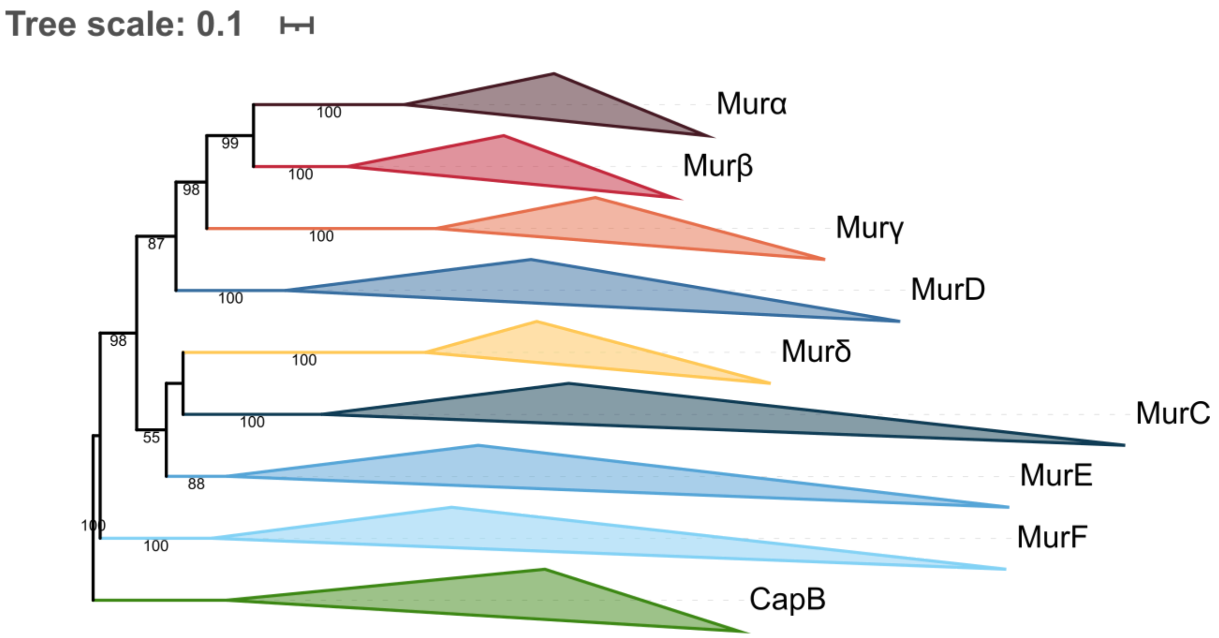


**Figure S21. Phylogenetic tree of the Mur domain-containing family rooted on CapB.** The tree was inferred from a matrix of **2519 sequences x 532 unambiguously aligned AAs** using IQ-TREE **under the C20+G4 model**. Tree visualization was performed using iTOL. Bootstrap values are shown if greater or equal to 50. Branches were collapsed on homogeneous sequence annotation.

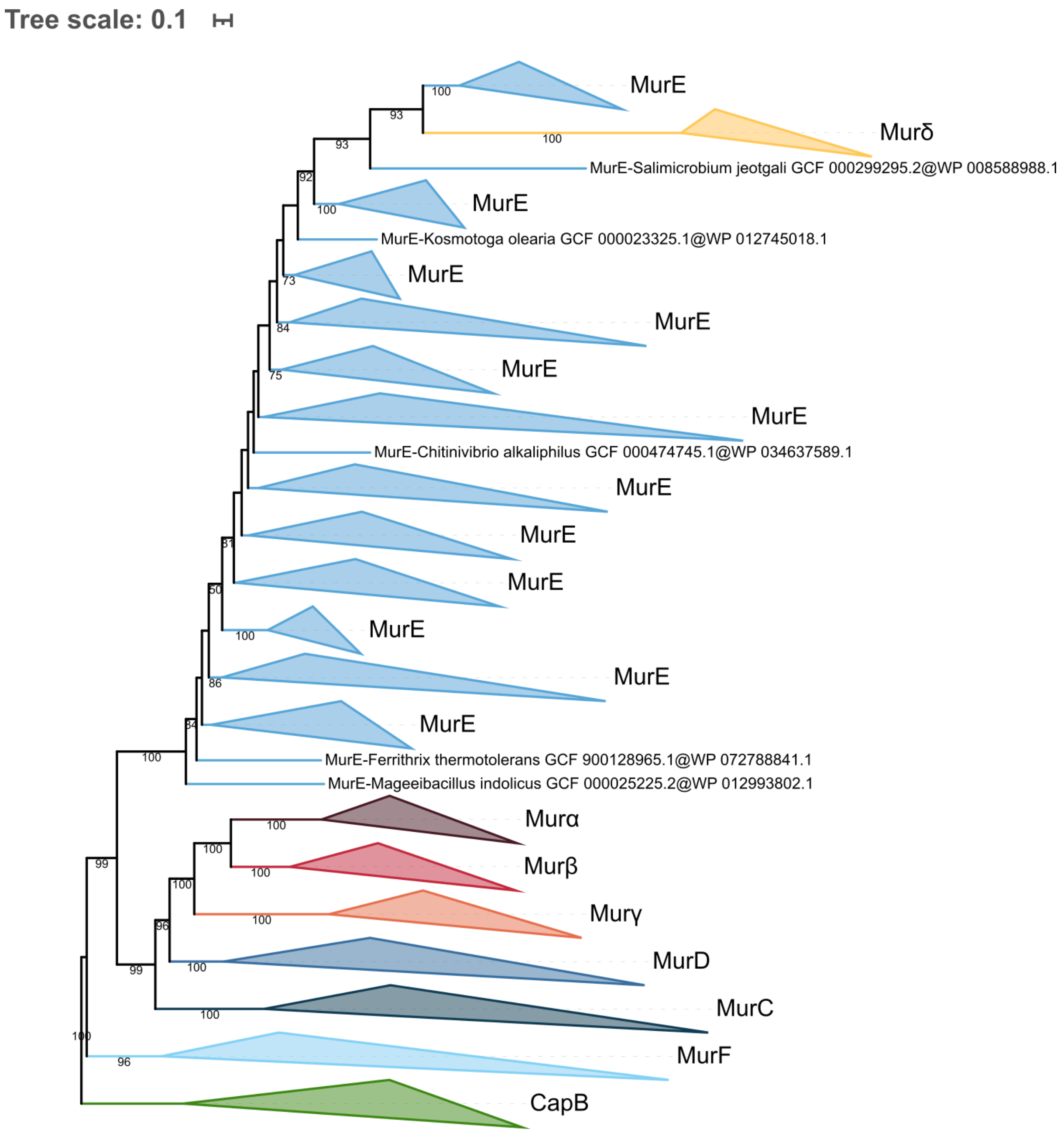


**Figure S22. Phylogenetic tree of the Mur domain-containing family rooted on CapB.** The tree was inferred from a matrix of **2519 sequences x 532 unambiguously aligned AAs** using IQ-TREE **under the C40+G4 model**. Tree visualization was performed using iTOL. Bootstrap values are shown if greater or equal to 50. Branches were collapsed on homogeneous sequence annotation.

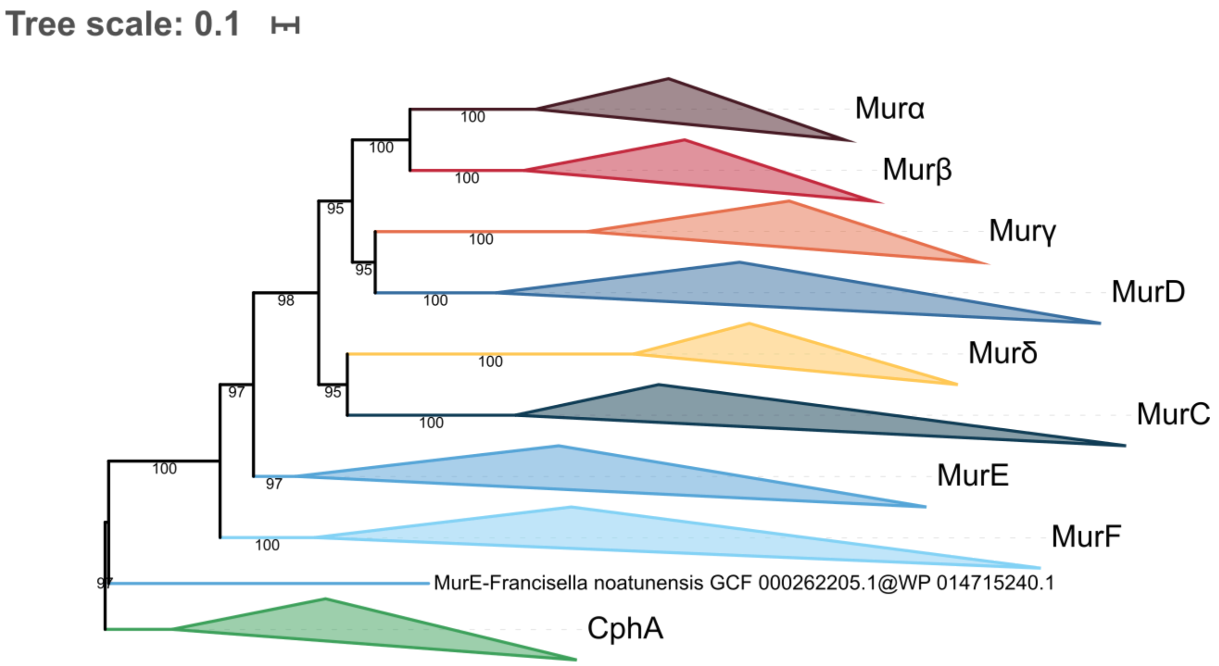


**Figure S23. Phylogenetic tree of the Mur domain-containing family rooted on CphA.** The tree was inferred from a matrix of **2461 sequences x 539 unambiguously aligned AAs** using IQ-TREE **under the LG4X+R4 model**. Tree visualization was performed using iTOL. Bootstrap values are shown if greater or equal to 50. Branches were collapsed on homogeneous sequence annotation.

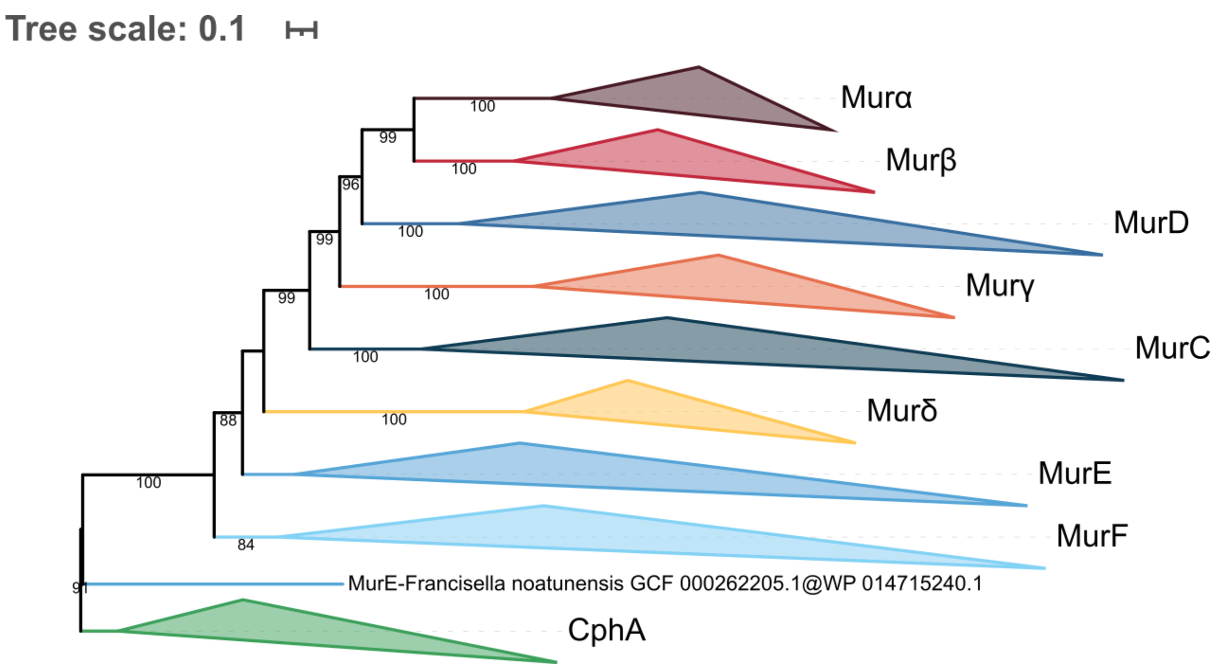


**Figure S24. Phylogenetic tree of the Mur domain-containing family rooted on CphA.** The tree was inferred from a matrix of **2461 sequences x 539 unambiguously aligned AAs** using IQ-TREE **under the C20+G4 model**. Tree visualization was performed using iTOL. Bootstrap values are shown if greater or equal to 50. Branches were collapsed on homogeneous sequence annotation.

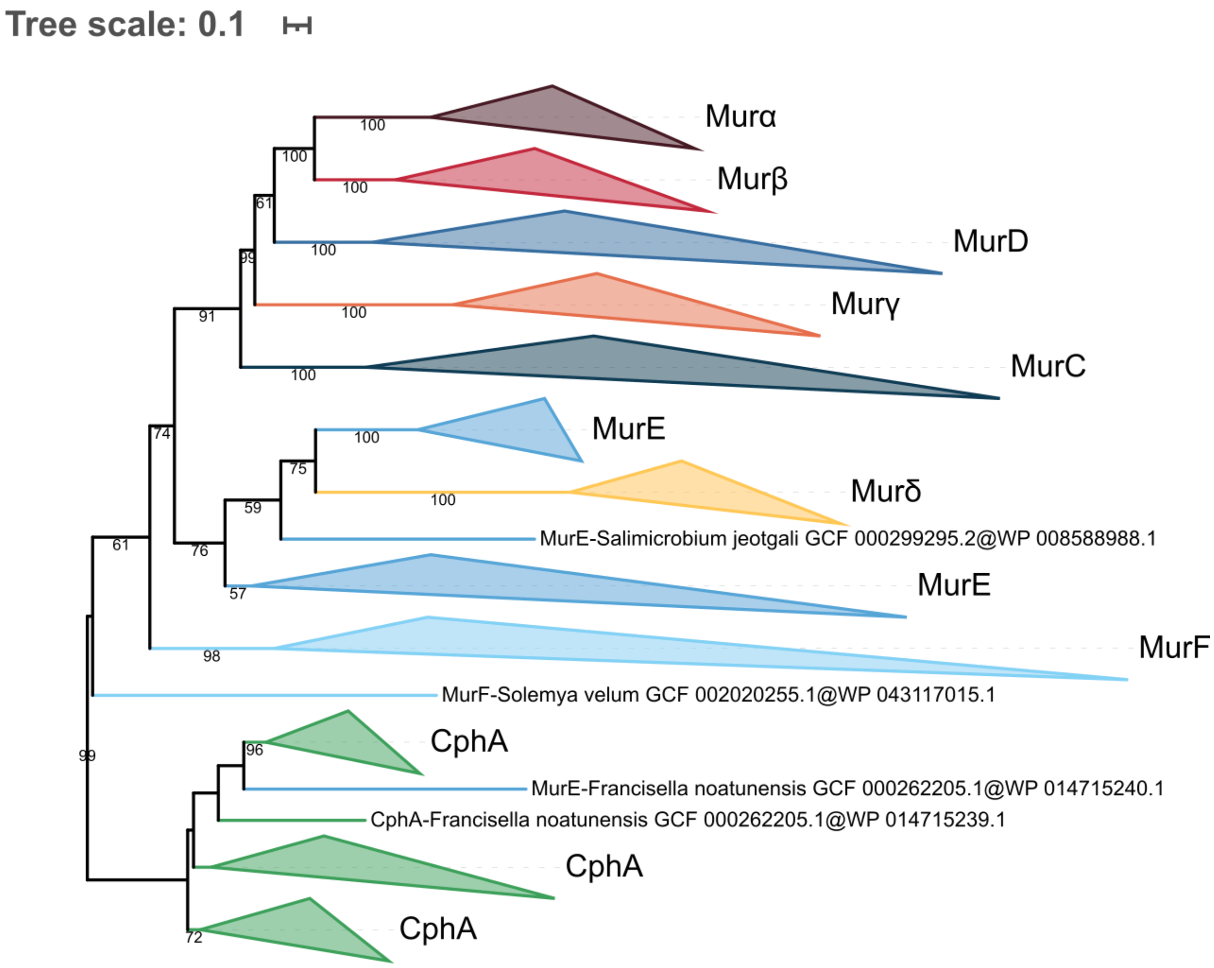


**Figure S25. Phylogenetic tree of the Mur domain-containing family rooted on CphA.** The tree was inferred from a matrix of **2461 sequences x 539 unambiguously aligned AAs** using IQ-TREE **under the C40+G4 model**. Tree visualization was performed using iTOL. Bootstrap values are shown if greater or equal to 50. Branches were collapsed on homogeneous sequence annotation.

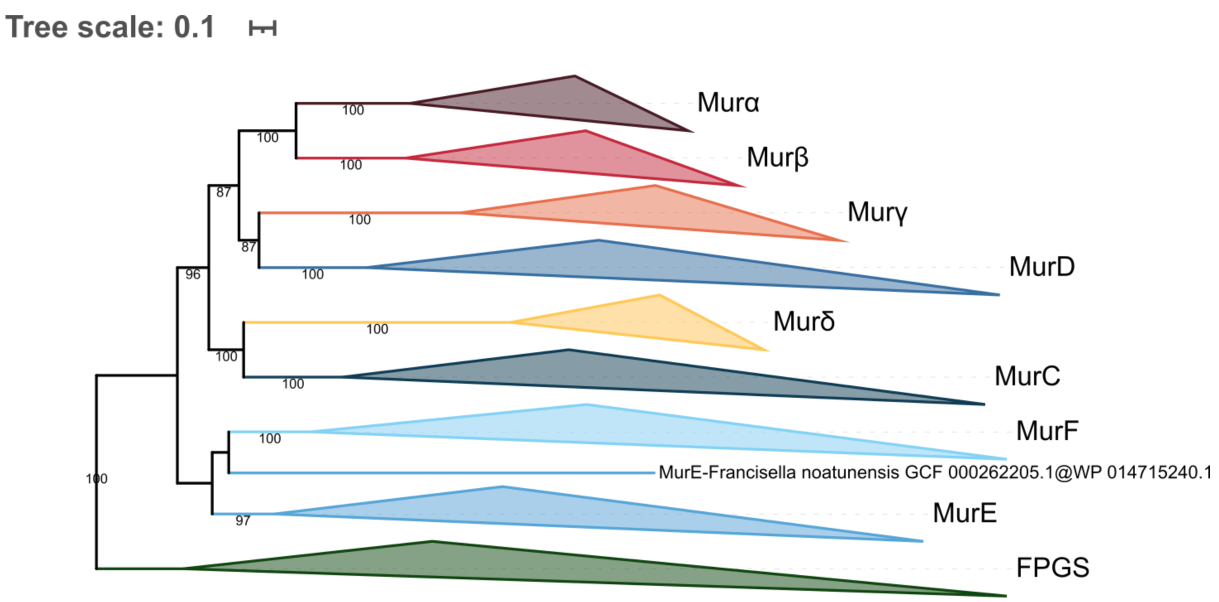


**Figure S26. Phylogenetic tree of the Mur domain-containing family rooted on FPGS.** The tree was inferred from a matrix of **3,046 sequences x 543 unambiguously aligned AAs** using IQ-TREE **under the LG4X+R4 model**. Tree visualization was performed using iTOL. Bootstrap values are shown if greater or equal to 50. Branches were collapsed on homogeneous sequence annotation.

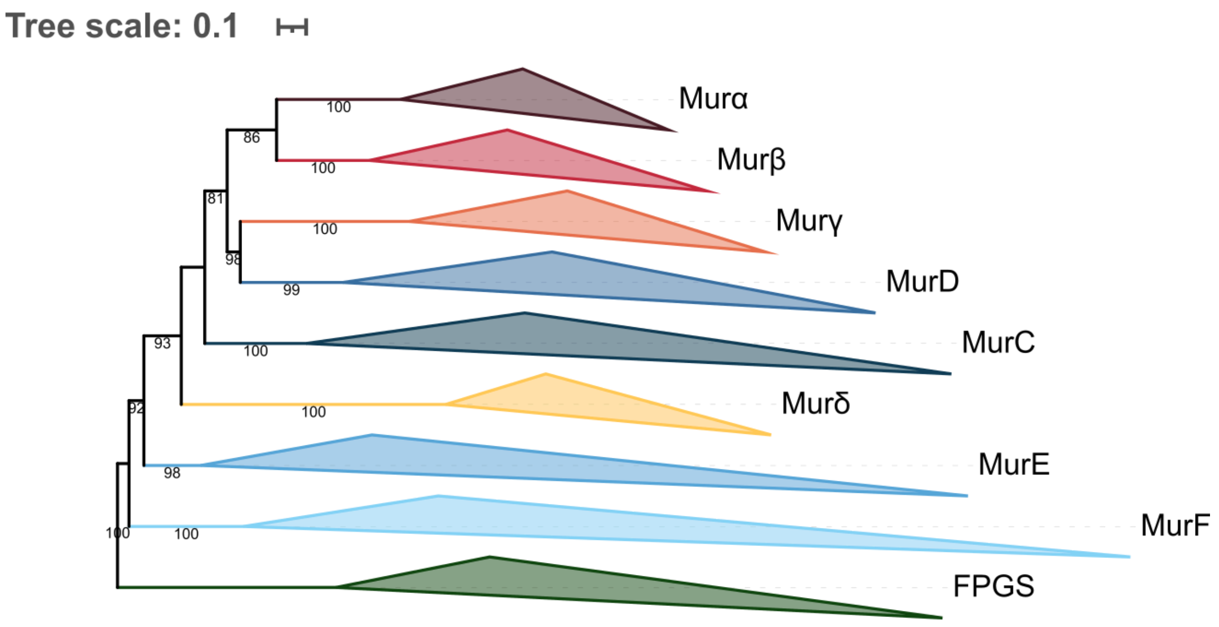


**Figure S27. Phylogenetic tree of the Mur domain-containing family rooted on FPGS.** The tree was inferred from a matrix of **3,046 sequences x 543 unambiguously aligned AAs** using IQ-TREE **under the C20+G4 model**. Tree visualization was performed using iTOL. Bootstrap values are shown if greater or equal to 50. Branches were collapsed on homogeneous sequence annotation.

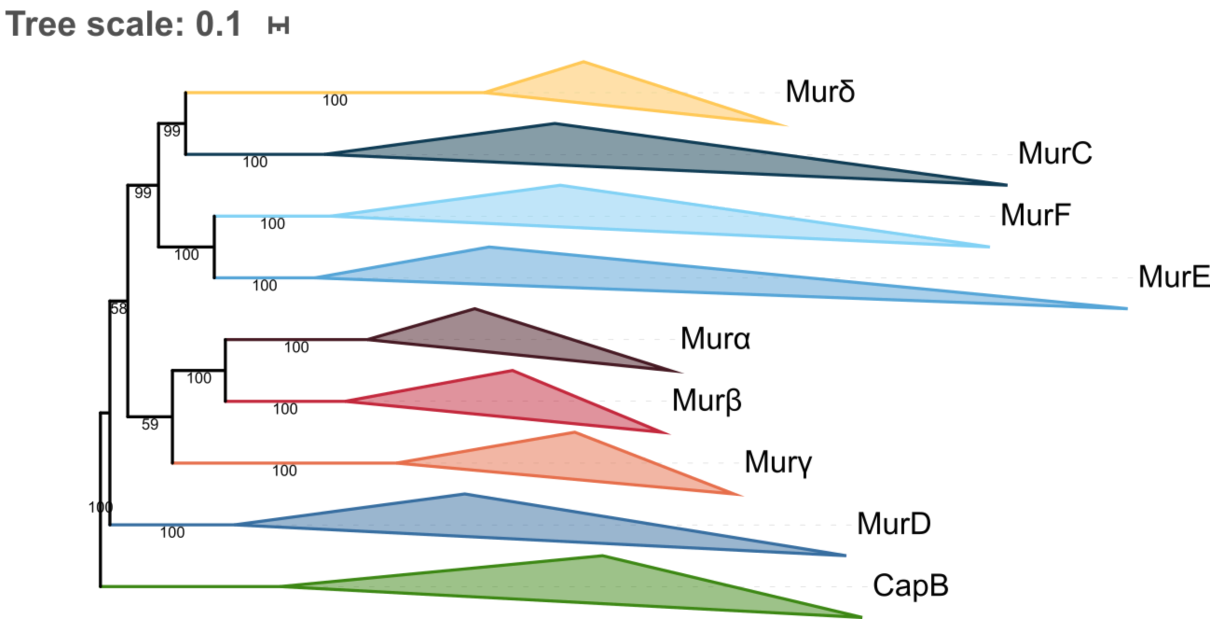


**Figure S28. Phylogenetic tree of the Mur domain-containing family rooted on CapB.** The tree was inferred from a matrix of **2519 sequences x 532 unambiguously aligned AAs** using IQ-TREE **under the PMSF LG+C60+G4 model** with 3000 iterations. The guide tree was the C40+G4 tree of Figure S22. Tree visualization was performed using iTOL. Bootstrap values are shown if greater or equal to 50. Branches were collapsed on homogeneous sequence annotation.

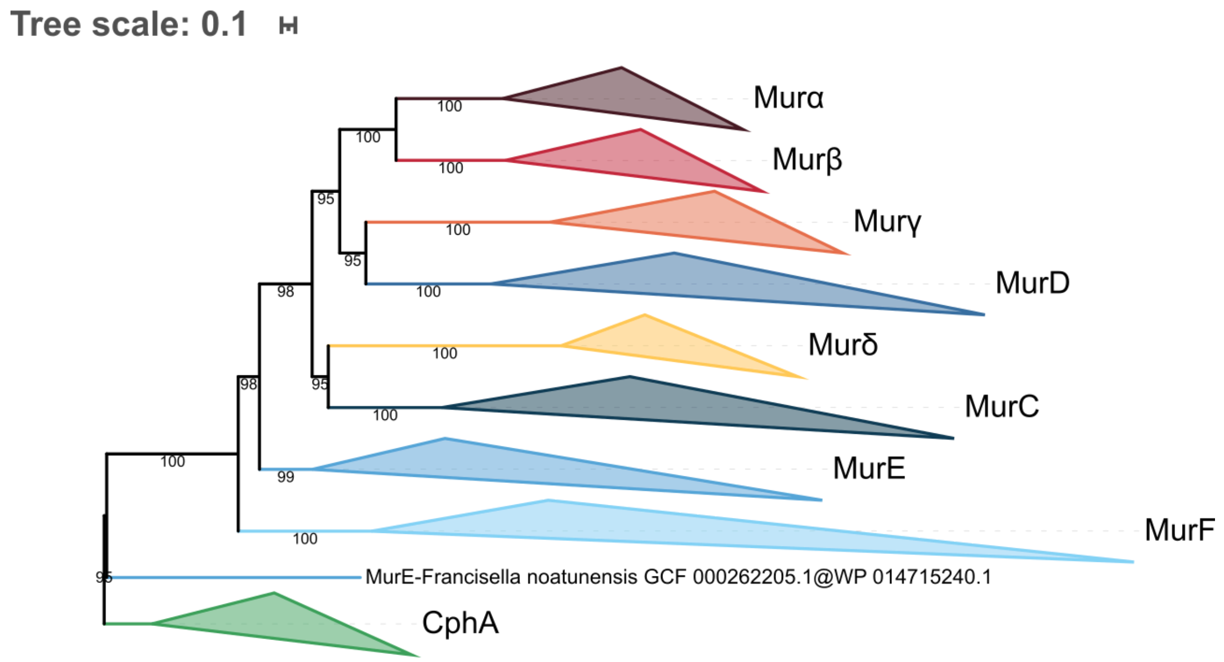


**Figure S29. Phylogenetic tree of the Mur domain-containing family rooted on CphA.** The tree was inferred from a matrix of **2461 sequences x 539 unambiguously aligned AAs** using IQ-TREE **under the PMSF LG+C60+G4 model** with 3000 iterations. The guide tree was the C40+G4 tree of Figure S25. Tree visualization was performed using iTOL. Bootstrap values are shown if greater or equal to 50. Branches were collapsed on homogeneous sequence annotation.

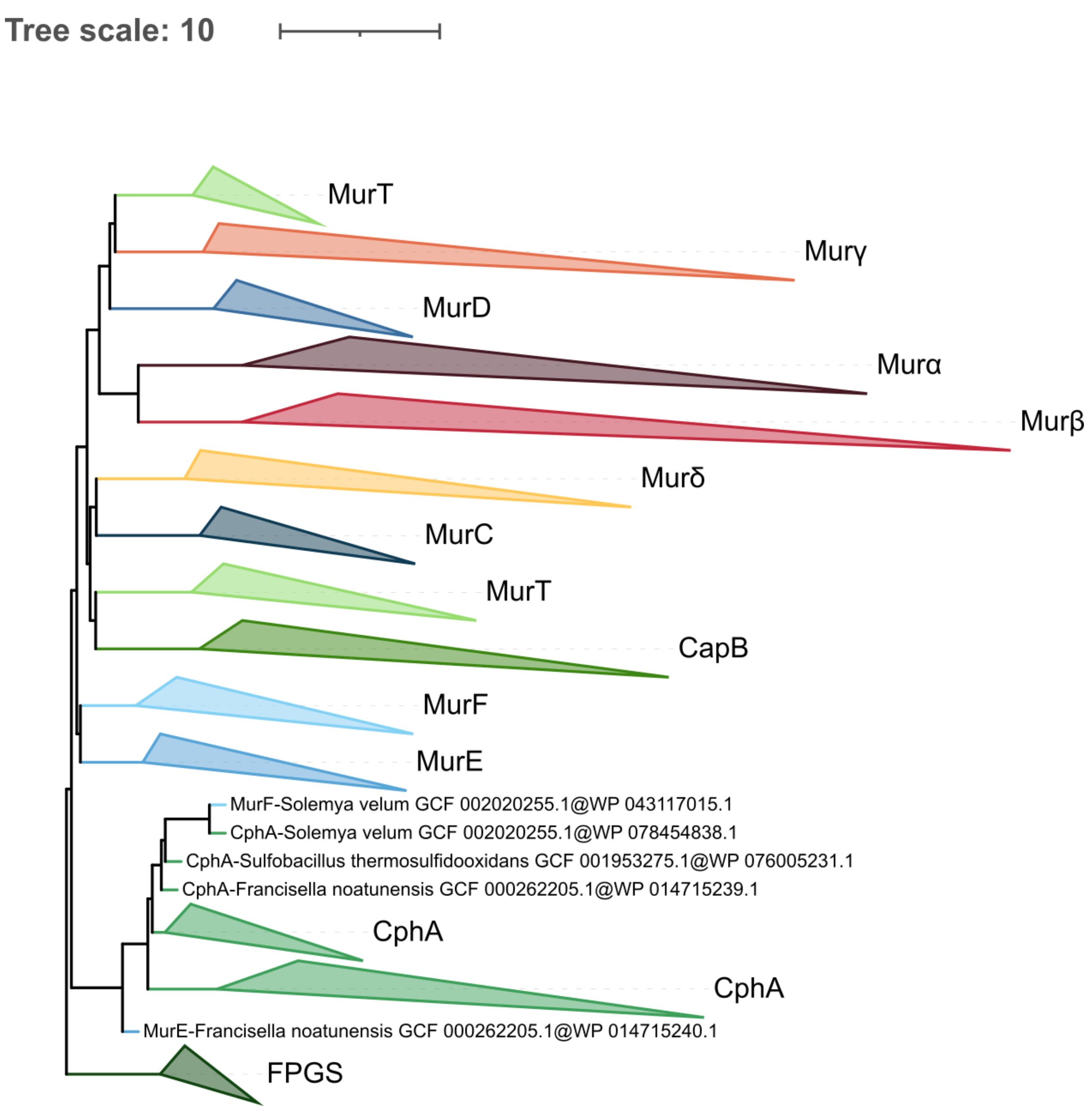


**Figure S30. Phylogenetic tree of the Mur domain-containing family rooted on FPGS. ASTRAL tree** inferred computed **from the 1000 LG4X+R4 species resampling trees**. Tree visualization was performed using iTOL. Branches were collapsed on homogeneous sequence annotation.

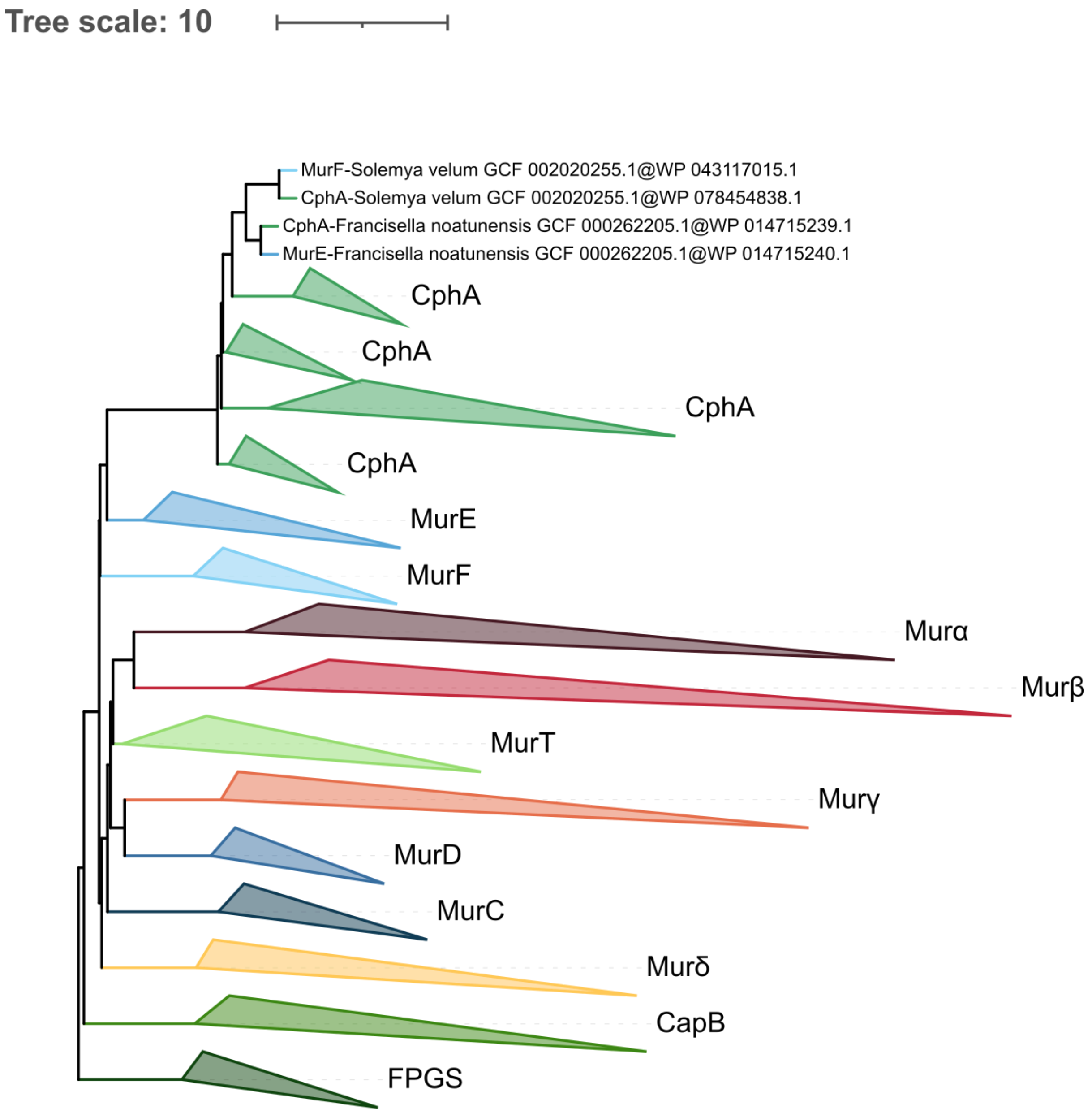


**Figure S31. Phylogenetic tree of the Mur domain-containing family rooted on FPGS. ASTRAL tree** computed **from the 1000 C20+G4 species resampling trees**. Tree visualization was performed using iTOL. Branches were collapsed on homogeneous sequence annotation.

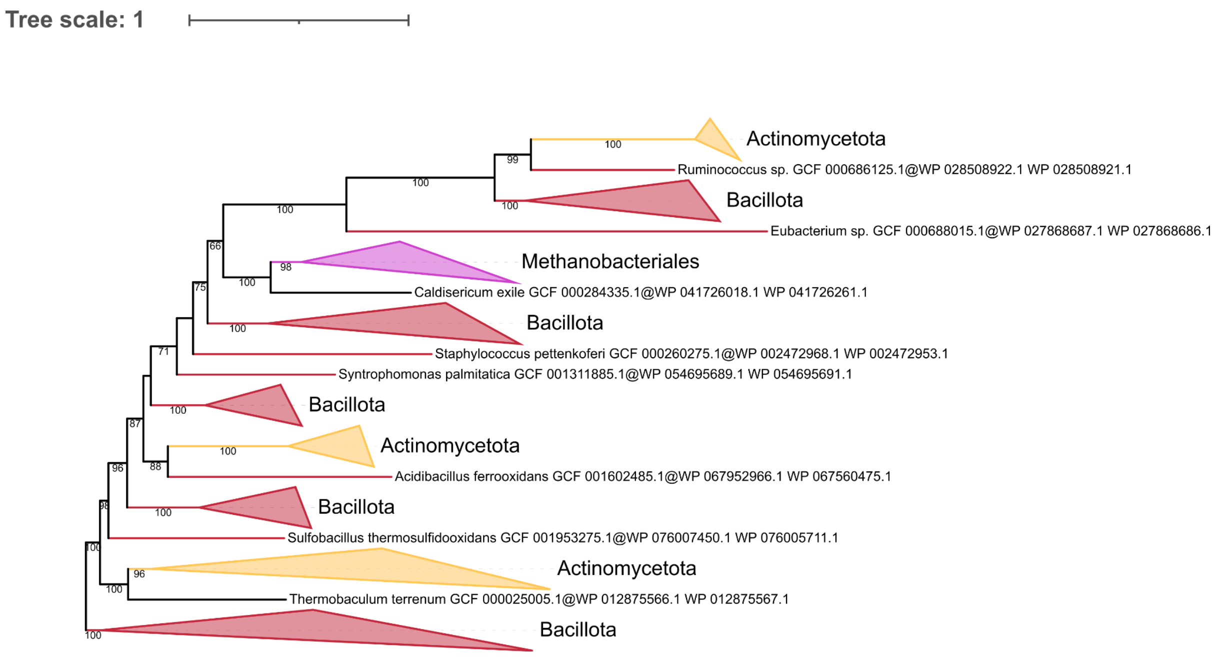


**Figure S32. Phylogenetic trees of the concatenation of MurT/GatD rooted on the largest Bacillota clan.** The tree was inferred from a matrix of 229 sequences x 653 unambiguously aligned AAs using IQ-TREE under the LG4X+R4 model. Tree visualization was performed using iTOL. Bootstrap support values are shown if greater or equal to 50. Branches were collapsed based on taxonomic clans.

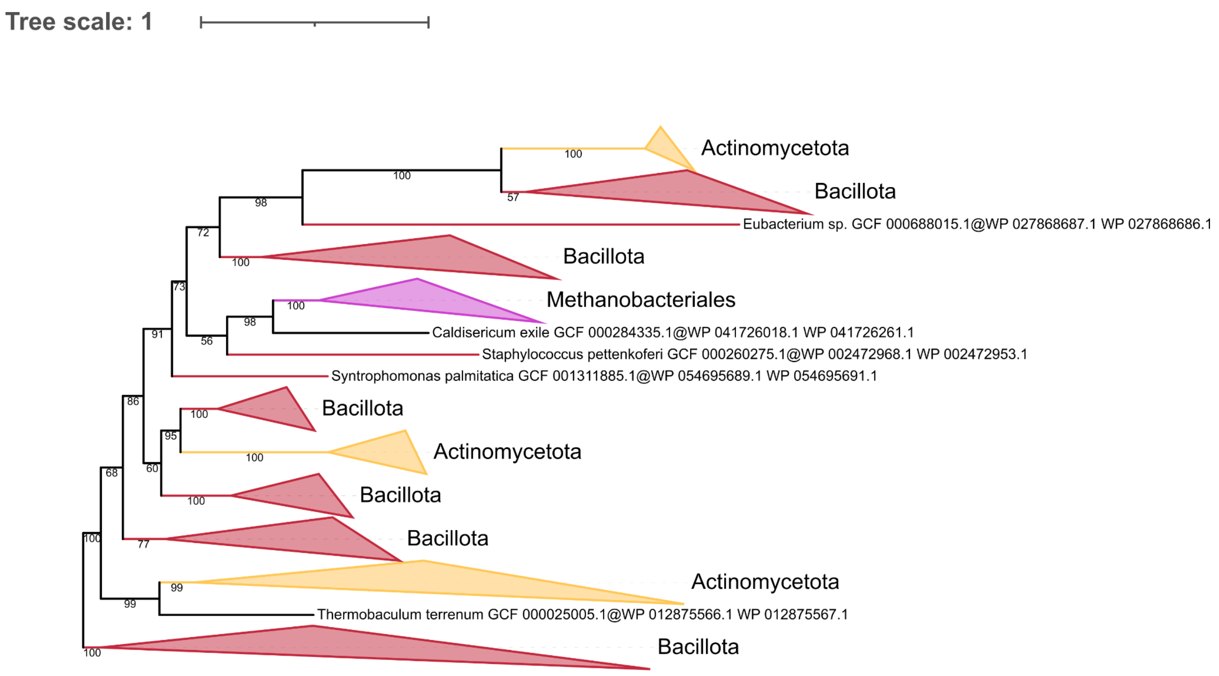


**Figure S33. Phylogenetic trees of the concatenation of MurT/GatD rooted on the largest Bacillota clan.** The tree was inferred from a matrix of 229 sequences x 653 unambiguously aligned AAs using IQ-TREE under the C20+G4 model. Tree visualization was performed using iTOL. Bootstrap support values are shown if greater or equal to 50. Branches were collapsed based on taxonomic clans.

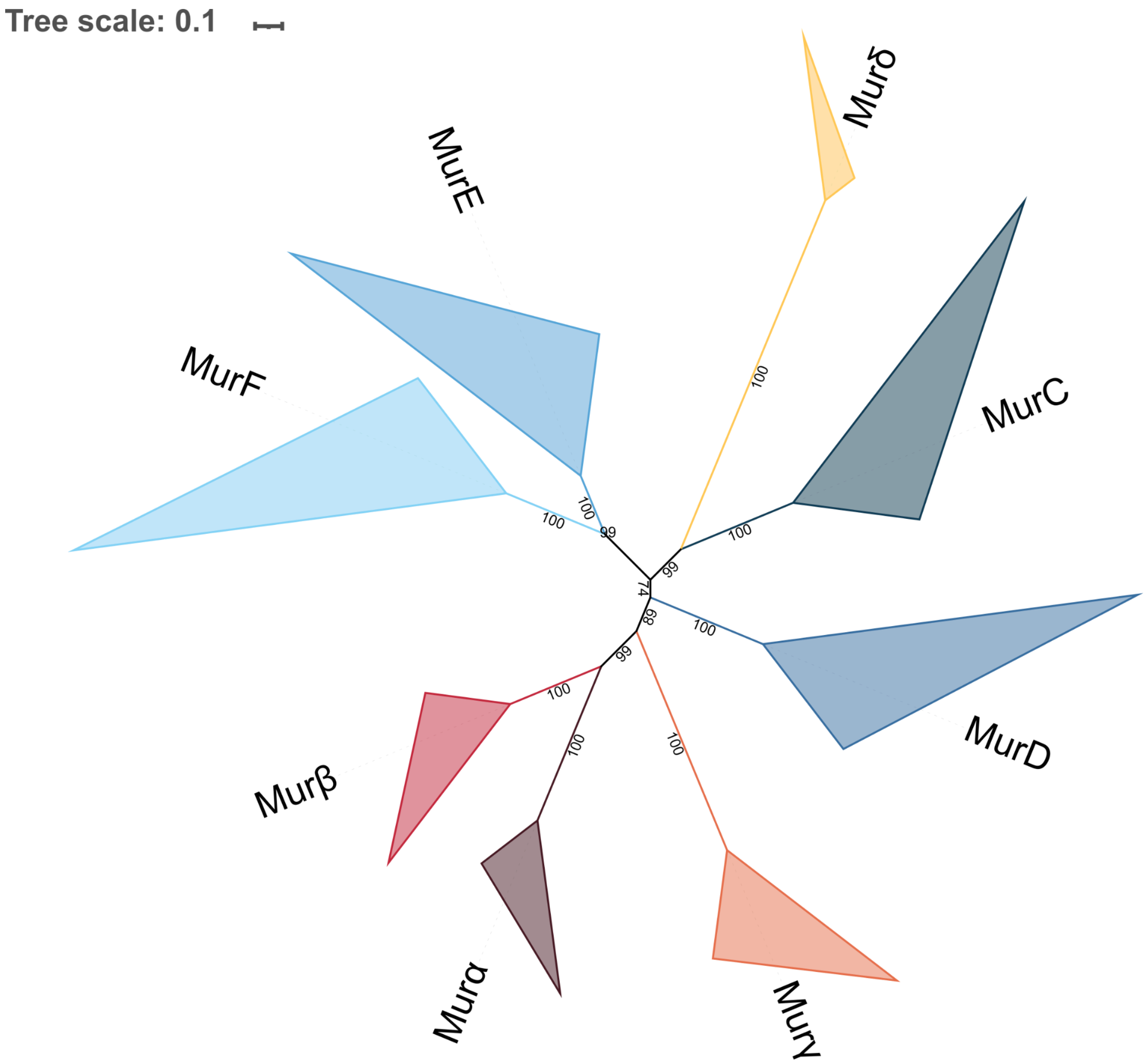


**Figure S34. Unrooted phylogenetic tree of the Mur domain-containing family.** The tree was inferred from a matrix of **2432 sequences x 528 unambiguously aligned AAs** using IQ-TREE **under the LG4X+R4 model**. Tree visualization was performed using iTOL. Bootstrap values are shown if greater or equal to 50. Branches were collapsed on homogeneous sequence annotation.

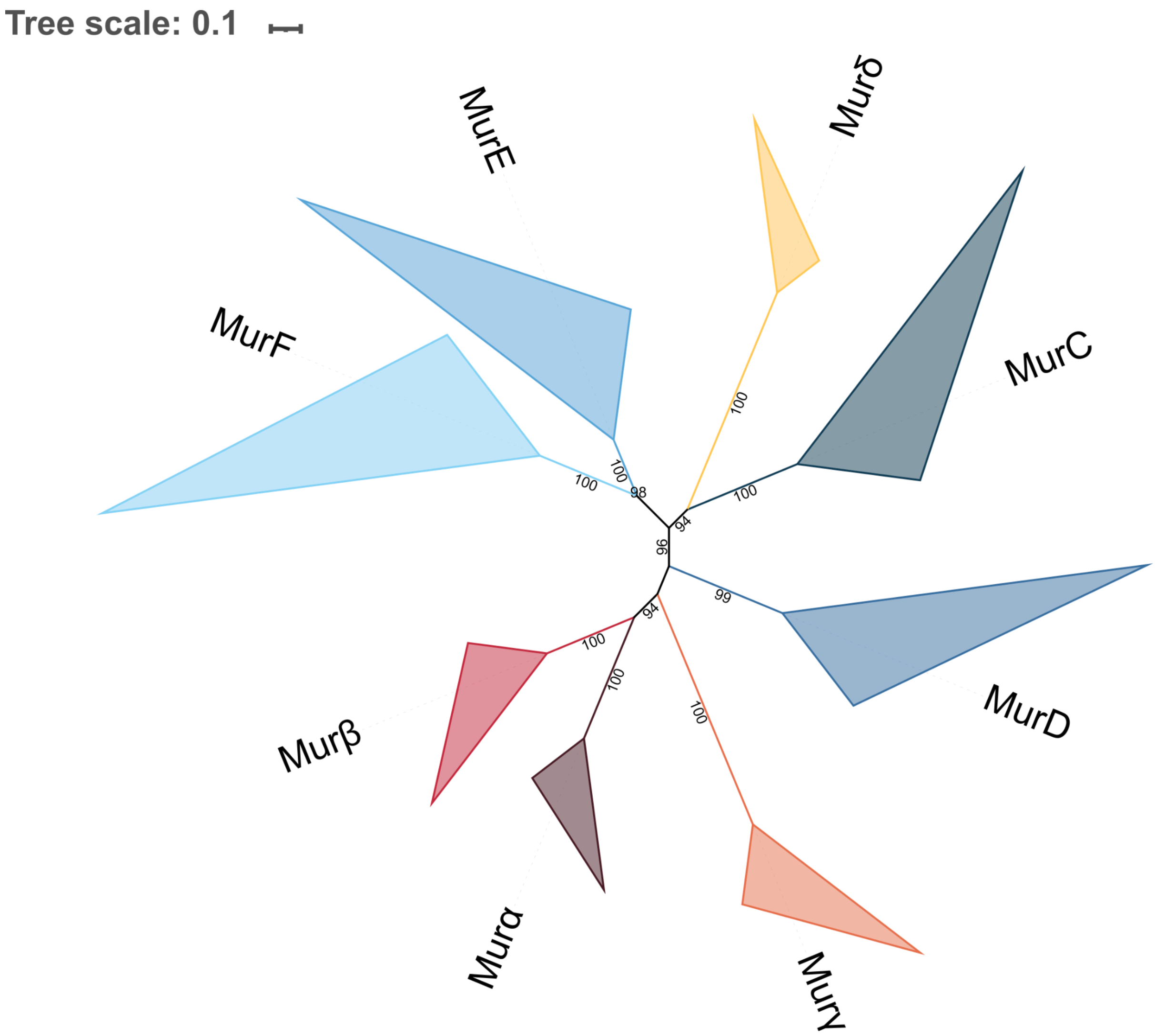


**Figure S35. Unrooted phylogenetic tree of the Mur domain-containing family.** The tree was inferred from a matrix of **2432 sequences x 528 unambiguously aligned AAs** using IQ-TREE **under the C20+G4 model**. Tree visualization was performed using iTOL. Bootstrap values are shown if greater or equal to 50. Branches were collapsed on homogeneous sequence annotation.

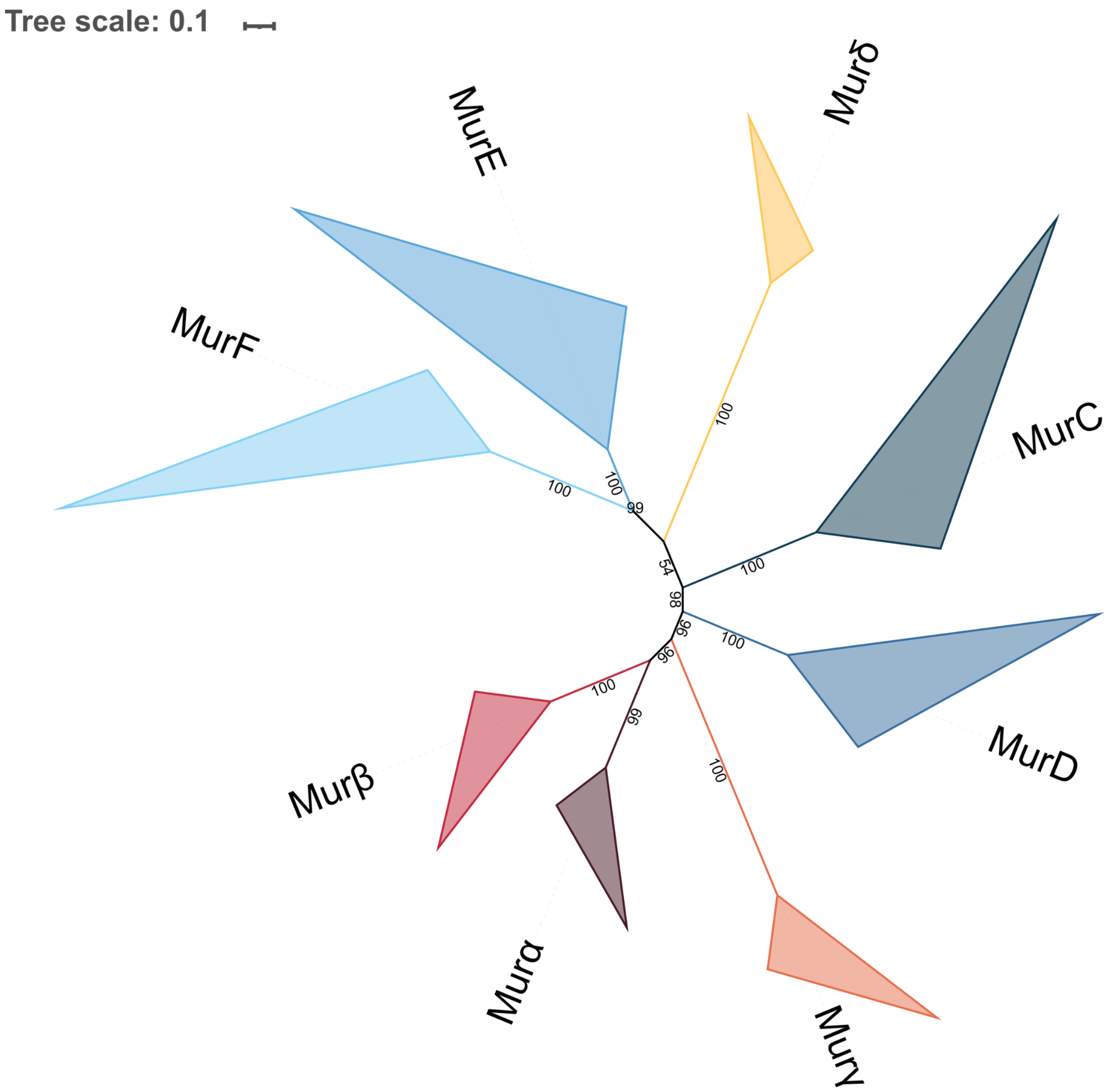


**Figure S36. Unrooted phylogenetic tree of the Mur domain-containing family.** The tree was inferred from a matrix of 2**432 sequences x 528 unambiguously aligned AAs** using IQ-TREE **under the C40+G4 model**. Tree visualization was performed using iTOL. Bootstrap values are shown if greater or equal to 50. Branches were collapsed on homogeneous sequence annotation.
