## Supplementary Tables legends for "Identification and Characterization of Archaeal Pseudomurein Biosynthesis Genes through Pangenomics"

**Table S1. Functional prediction of InterProScan along with the prediction of signal peptide type and the number of predicted transmembrane segments.** Sheets 1 and 2 include the Orthologous Groups (OGs) identified using taxonomic filters, sheet 3 includes the OGs identified through synteny, sheet 4 includes OGs identified using the three HMM profiles from the CDD database and sheet 6 includes OGs from the regulon analysis (see supplemental data).

**Table S2. Detailed distribution patterns of the 49 retained orthologous groups (OGs) using either the bacterial database or the larger prokaryotic database.**

**Table S3. Jackknife support values computed from the 1000 replicates of species resampling under three phylogenetic models: LG4X+R4, C20+G4 and C40+G4.** The second table shows jackknife support values when considering two divergent sequences, one annotated as a MurE and the other as a MurF, as CphA, while the third table further considers the clan of MurT-like.

**Table S4. Different combinations of intergenic regions.** Lowercase letters refer to as the intergenic region indicated in Figure S2, while uppercase letters correspond to one of the five PM-containing archaea: MC = *Methanobacterium congolense*; MS = *Methanobrevibacter smithii*; M = *Methanopyrus* sp.; MT = *Methanothermobacter thermautotrophicus*; MF = *Methanothermus fervidus*. Bold letters correspond to intergenic regions actually used by MEME to discover a motif.
